# Chain entropy modulates cooperativity selectively within intermediate sub-populations during protein unfolding

**DOI:** 10.64898/2026.03.08.710427

**Authors:** Anushka Kaushik, Jayant B. Udgaonkar

**Author notes:** Corresponding author: Jayant Udgaonkar.

## Abstract

Protein unfolding invariably appears to be a cooperative transition; yet, the molecular basis by which structural elements could unfold in a coordinated manner remains unresolved. Here, the unfolding mechanism of the naturally occurring heterodimeric protein double-chain monellin (dcMN) was characterized using site-specific time-resolved FRET and fluorescence anisotropy decay measurements made under equilibrium conditions. Although ensemble-averaged measurements suggested an apparently cooperative transition, population-level analysis using the maximum entropy method coupled to time-resolved FRET revealed pronounced conformational heterogeneity, with partially contracted (N-like) coexisting with partially expanded (U-like) sub-populations during unfolding. Time-resolved fluorescence anisotropy decay measurements independently demonstrated that local motional constraints are lost gradually and asynchronously across different regions of the protein. The N-like sub-populations underwent cooperative expansion across both intra- and inter-chain segments, indicating coordinated responses when inter-chain coupling is maintained. In contrast, the U-like sub-populations displayed pronounced chain-specific, non-cooperative behavior, consistent with independent unfolding of the two chains following loss of coupling. Comparison with a covalently linked single-chain variant demonstrates that chain connectivity suppresses heterogeneity and enforces coordinated unfolding. These results identify restriction of chain entropy arising from inter-chain coupling and covalent connectivity as a molecular determinant that governs whether heterogeneous intermediate sub-populations unfold cooperatively or in a chain-specific manner.

## Introduction

The extent and origin of cooperativity in protein folding and unfolding reactions are not yet fully understood. Folding transitions are often described as “two-state” when monitored by ensemble-averaging probes, which typically report sigmoidal transitions between the native (N) and unfolded (U) states.^1–3^ This seeming all-or-none behaviour led to the view that proteins have evolved to fold and unfold cooperatively, so that the population of partially folded intermediates that could be aggregation-prone, is minimal.^4–5^ A two-state description, however, does not exclude the possibility that high-energy intermediates are populated transiently during folding. When folding is probed by more sensitive techniques, partially folded intermediates have been detected and structurally characterized, and in several cases intermediates stable enough to accumulate to measurable extents, have been identified.^6–13^ Indeed, even when the thermodynamic and kinetic criteria for two-state folding are satisfied, partially unfolded intermediates may still be sparsely populated, and different structural segments may lose structure gradually or asynchronously.

A true two-state conformational change in a protein would be one in which different structural regions undergo synchronous change. But unfolding under equilibrium conditions can appear to be two-state even when (i) different structural regions of the protein have distinct local stabilities that are distributed around the global stability, resulting in unfolding being heterogeneous and not fully cooperative,^14–19^ and (ii) certain regions undergo gradual unfolding, in which structural order is lost progressively.^8, 20–28^ In such cases, the non-cooperative behaviour of some structural elements may remain hidden beneath the apparent global cooperativity reported by ensemble-averaging probes, and be revealed only by site-specific, high-resolution methods such as single-molecule fluorescence resonance energy transfer (sm-FRET),^20^ time-resolved FRET,^12, 29–31^ and hydrogen exchange coupled to mass spectrometry (HX-MS).^11, 22, 32–33^ For several proteins, these methodologies have revealed that (un)folding can be slowed down not just by a dominant high free energy barrier, but also by a rugged landscape with many distributed small free energy barriers,^34–36^ leading to gradual, asynchronous structural transitions.^8, 12, 21, 37–39^ What remains unclear is whether deviations from two-state behaviour originate solely within secondary structural elements or whether they also involve long-range interactions. How the intrinsic differences in cooperativity of formation of different structural motifs are influenced by their placement within the overall protein architecture remains poorly understood. Moreover, whether multi-chain proteins display distinctive cooperative or non-cooperative unfolding patterns compared to their single-chain counterparts is an open question.

The small protein monellin is an ideal model system for investigating folding cooperativity, as it exists in two closely related forms: a naturally occurring heterodimer (dcMN),^40–41^ and an artificially created single-chain variant (MNEI)^42–43^ in which the C-terminus of chain B (in β2) is linked covalently to the N-terminus of chain A (in β3) *via* a short Gly-Phe peptide linker (Figure 1a). The native-state structures of dcMN and MNEI are nearly identical^41, 44^ except at the linker region (Figure 1a). These two variants of monellin offer a unique opportunity to directly compare the unfolding cooperativity of single-chain and double-chain architectures, while minimizing confounding differences in sequence or structure, thereby isolating the effect of chain connectivity.

**Figure 1.**
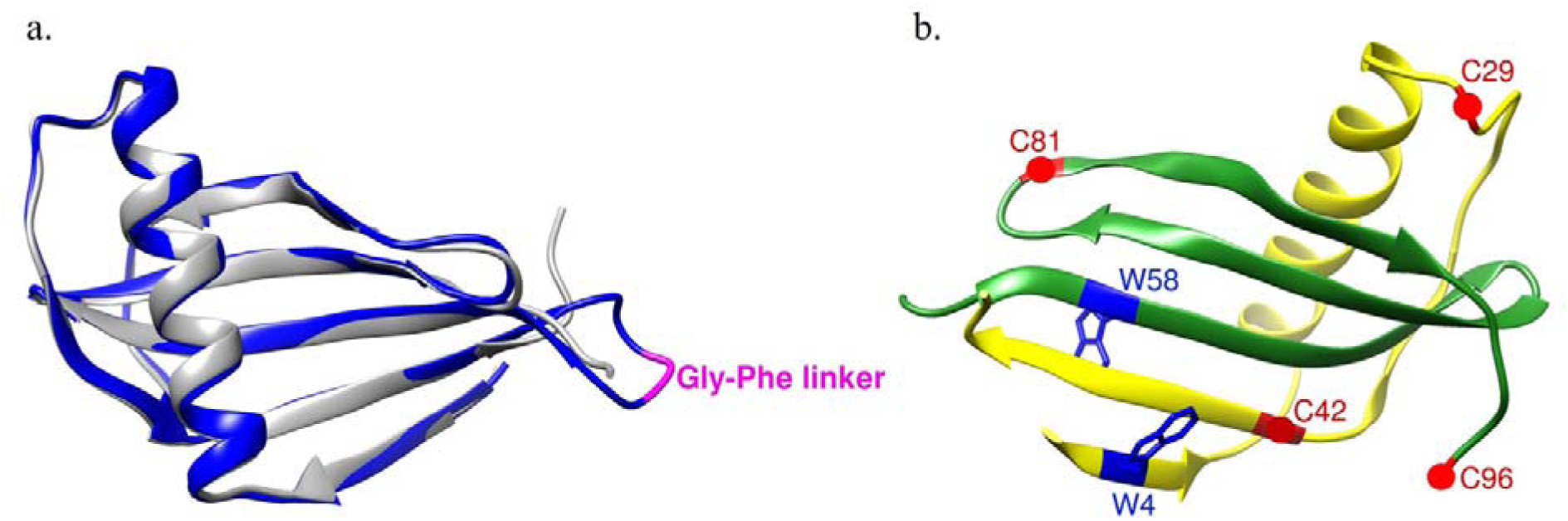
a) Structural comparison of MNEI and dcMN. The Gly–Phe linker (shown in pink) joins the two chains of dcMN (shown in grey) to yield MNEI (shown in blue). b) Structure of dcMN showing the different residues used for monitoring FRET. Chain B of dcMN is shown in yellow, while chain A is shown in green. Wild-type dcMN has one Trp residue (W4) and one Cys residue (C42). The side chains of the Trp residues (used as the FRET donor) are shown as blue rings, and the Cys residues (to which the FRET acceptor TNB was attached) are shown as red spheres. FRET was measured in single tryptophan-containing, single cysteine-containing variants. FRET changes were monitored within chain B (in W4C42), within chain A (in W58C81), between the two chains (in W4C96), and across the helix (in W4C29) present in chain B. The distances between the centre of the Trp ring and Cβ atoms of the Cys residue (attached to the acceptor) in the N state, were 13.0 Å (W4C42), 14.3 Å (W58C81), 16.0 Å (W4C96) and 22.5 Å (W4C29). The structure has been drawn using the Chimera software and the PDB IDs 1IV7 and 3MON.

It is known that MNEI exhibits deviations from two-state behaviour: kinetic and equilibrium studies using multisite tr-FRET and HX-MS have revealed continuous, segmental expansion of the β-sheet during unfolding^10, 21^ and limited cooperativity. ^11–12, 22^ Other studies have shown asynchronous folding of the helix,^45^ and the transient population of metastable intermediates^13, 39, 46^ that further underscore the heterogeneous and non-cooperative nature of the folding and unfolding pathways. dcMN appears to differ from MNEI in its stability^33, 41^ as well as in its folding cooperativity,^22, 33, 47^ and it is not known whether these differences arise because MNEI is an artificially created protein. An HX-MS study on dcMN^33^ showed that unfolding under native and mildly denaturing conditions involves three non-cooperative kinetic phases followed by a slower cooperative phase. Interestingly, while only β3 in chain A unfolds cooperatively under strongly native conditions (≤0.4 M GdnHCl), both β3 and β2 in chain B unfold cooperatively under mildly denaturing concentrations (0.6–1 M GdnHCl). However, HX-based methods probe only those regions that display measurable protection, and typically provide little or no information on more than half of the structure, because that offers little or no protection against HX. As a result, intrinsically flexible segments and transiently unprotected regions, which may include parts of the inter-chain dimer interface, are often not captured, even though they may play a key role in coordinating conformational transitions during unfolding.

To understand how protein architecture and chain connectivity modulate folding cooperativity and heterogeneity, MEM-coupled time-resolved FRET measurements were used together with site-specific fluorescence anisotropy measurements to study the unfolding of dcMN under equilibrium conditions. Four FRET pairs were designed: two intra-chain pairs to map distance distributions within the structural cores of chain A (in W58C81) and chain B (in W4C42), one inter-chain pair (in W4C96) to monitor the distance distribution across the two chains, and another (in W4C29) to map the helix segment (Figure 1b). Measurements of the time-resolved fluorescence anisotropy decay of Trp58 in W58C81 provide complementary insights into the motional dynamics at the dimer interface and within chain A, while those of Trp4 in W4C42 report on chain B. These approaches together reveal localized deviations from two-state behaviour that remain hidden when ensemble-averaging probes are utilized to study unfolding. By following the GdnHCl concentration-dependent evolution of MEM-derived lifetime distributions and anisotropy-derived dynamics across intra-chain cores, inter-chain contacts, the helix, and the dimer interface, the structural basis of unfolding cooperativity in dcMN has been delineated.

## Results

### The structure and stability of the protein were not significantly perturbed by mutation and labeling

To monitor distinct intramolecular distances in dcMN, four single-Trp, single-Cys variants were generated (see Materials and Methods). The tryptophan residues (Trp4 in chain B and Trp58 in chain A) served as FRET donors, and a TNB group covalently attached to a unique cysteine acted as the acceptor. dcMN contains a native Cys residue at residue position 42, and a cysteine residue was introduced at residue position 29, 81, or 96 in place of Cys42 to enable site-specific labelling (see Materials and Methods). The unfolding transitions of these protein variants were then monitored by FRET at various concentrations of GdnHCl. The fluorescence emission maximum of both Trp4 in chain B (for W4C42, W4C96 and W4C29) and Trp58 in chain A (for W58C81), upon excitation at 295 nm, was at ∼ 347 nm for the native unlabeled protein (Figure S1) indicating that the local environments of Trp4 and Trp58 were similar. Upon unfolding, the wavelength of maximum fluorescence emission increased to 356 nm for all the protein variants, reflecting full exposure of W4 or W58 to solvent. Upon labeling with the TNB acceptor, quenching of the Trp fluorescence intensity was observed at all wavelengths, and the extent of quenching was different for the N and U states. The extent of quenching upon TNB-labeling was different for the different protein variants, indicating that quenching was distance-dependent, and hence, due to FRET. The extent of quenching was also found to be independent of protein concentration, ruling out intermolecular energy transfer (data not shown). The secondary structure, as can be seen in the far-UV CD spectra (Figure S2) was not perturbed significantly by mutation and labeling. The minor differences in the spectra of the variants can be attributed to differences in the contributions of Trp and Tyr at residue positions 4 and 58.^48–49^

The stabilities of all the unlabeled proteins were not significantly different (< 1 kcal mol^-1^) from that of wt dcMN (W4C42) (Figure 2 and Table S1), indicating that the mutations caused minimal perturbation. Labeling with TNB also had little effect on stability for the W58C81, W4C96, and W4C29 variants, but the W4C42 variant was slightly destabilized. Cys42 is completely buried in the N state, and the covalent modification of this residue by TNB, may have disrupted packing interactions and thereby caused a minor decrease in stability. Similar destabilization upon TNB-labeling of Cys42 had also been observed for MNEI.^12^ Nevertheless, the overall folding and unfolding properties were not altered significantly: the refolding kinetics of the labeled and unlabeled variants were indistinguishable from those of wt dcMN (data not shown).

**Figure 2.**
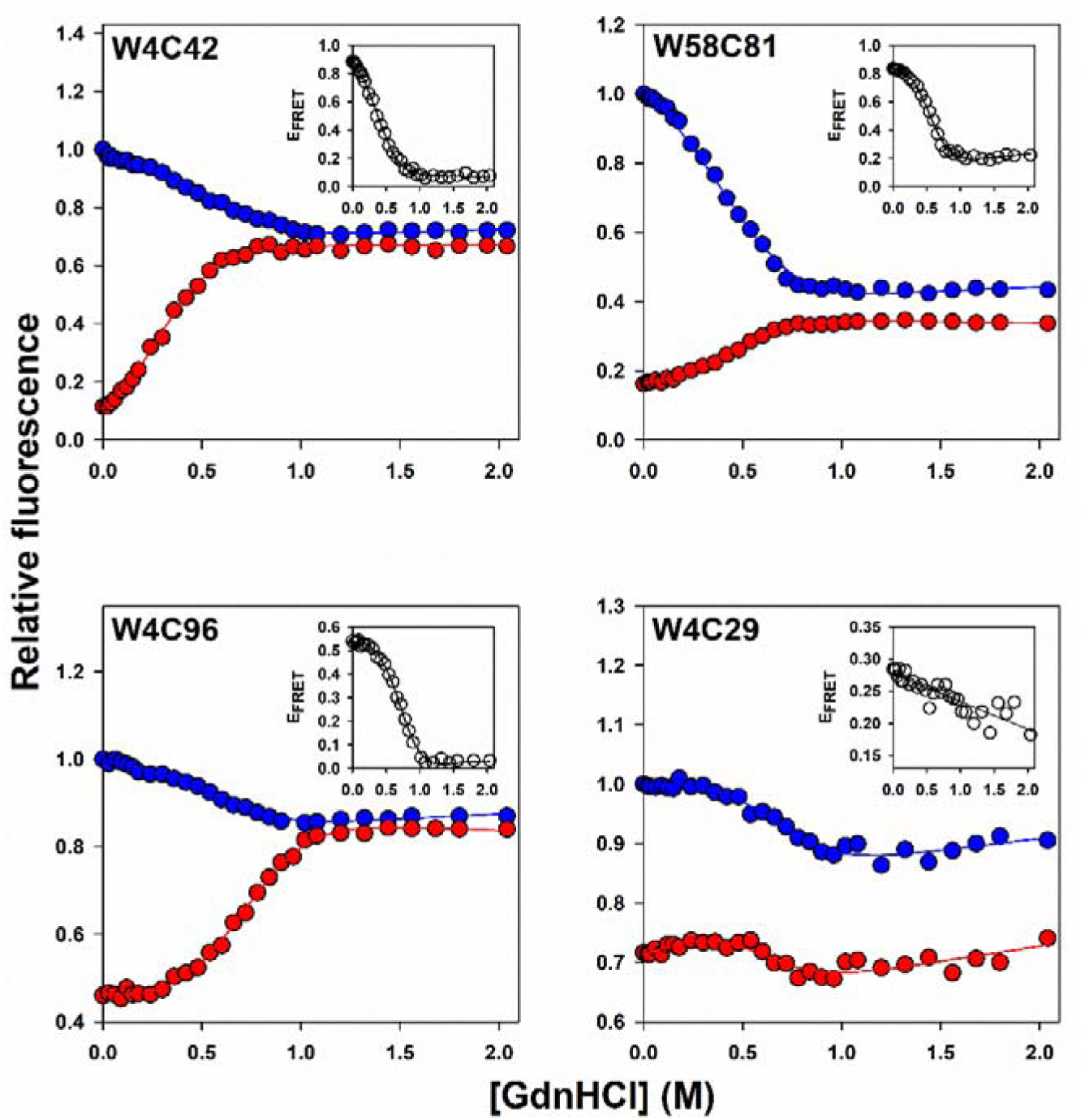
Equilibrium unfolding transitions of different variants of dcMN monitored by steady-state FRET measurements. The blue and red circles correspond to the fluorescence intensities of the unlabeled and TNB-labeled proteins, respectively. The solid line passing through each dataset is a non-linear, least-squares fit to a two-state unfolding model for a heterodimeric protein.^41^ The thermodynamic parameters obtained are listed in Table S1. The inset in each panel shows the dependence of the FRET efficiency on GdnHCl concentration. The solid line shown in the inset of the panel for W4C29 is drawn to guide the eye.

### Dependence of FRET efficiency on GdnHCl concentration appeared non-sigmoidal for the helix segment

The equilibrium unfolding transitions of W4C42, W58C81, W49C96 and W4C29 monitored by fluorescence, appeared to be two-state in nature, as indicated by the sigmoidal dependence of fluorescence intensity on GdnHCl concentration (Figure 2). The dependence of FRET efficiency on GdnHCl concentration (insets, Figure 2) also appeared to be describable as a two-state transition, for all variants except W4C29, which showed a broad non-sigmoidal transition.

The dependence of the mean lifetime on GdnHCl concentration, derived from the discrete analysis of time-resolved fluorescence lifetime decay curves obtained at different GdnHCl concentrations, compared very well with that monitored by Trp fluorescence (steady-state measurements) (Figure S3). Moreover, for all the protein variants, there was good agreement between the FRET efficiencies obtained from fluorescence intensity and lifetime measurements (Figures 2 and 3).

**Figure 3.**
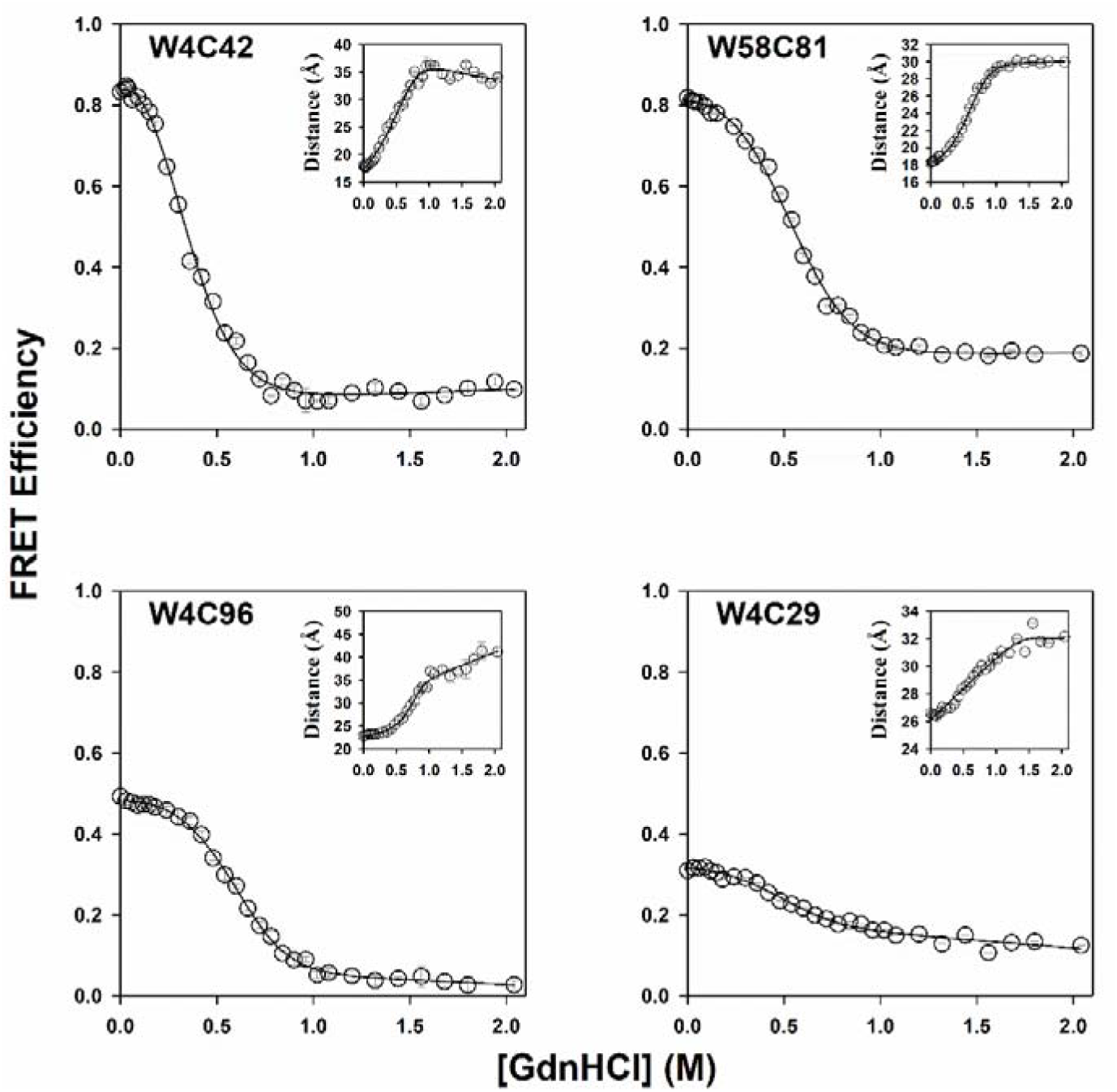
Equilibrium unfolding transitions of different variants of dcMN monitored by time-resolved FRET measurements. FRET efficiency was calculated using mean lifetimes that were obtained from discrete analysis of the time-resolved data for both the unlabeled and the TNB-labeled protein variants. The inset in each panel shows the dependence of the donor–acceptor distances on GdnHCl concentration. The distances were determined using R_o_ values (Table S2; Materials and Methods). The error bars represent the spread in the data obtained from two independent experiments.

However, based on these measurements, the equilibrium unfolding transitions could not be classified as either strictly cooperative or gradual, as these measurements could not resolve distinct conformational subpopulations populated at different GdnHCl concentrations. It should be noted that negligible FRET is observed for the segment mapping the distance across the two chains (W4C96) at GdnHCl concentrations above 1.5 M (Figures 2 and 3). If this FRET amplitude was nevertheless used to infer the distance separating Trp4 and Cys96-TNB, a distance > 40 Å was obtained, which exceeded the reliable distance-sensitive range of the FRET pair (R[ = 22.7 Å). These distances should therefore be regarded as qualitative indicators of chain separation rather than precise quantitative measurements. In addition, far-UV CD spectra for the segments mapping the cores of the two chains (in W4C42 and W58C81), as well as for the inter-chain segment (in W4C96), showed a complete loss of secondary structure at 2 M GdnHCl (Figure S2). Together, these observations indicated that at high denaturant concentrations, the two chains have separated from each other.

### MEM analysis revealed the coexistence of cooperative and continuous unfolding

Fluorescence intensity decay curves were analyzed using the Maximum Entropy Method (MEM) to obtain fluorescence lifetime distributions (Figure 4). In the case of W4C42-TNB, W58C81-TNB and W4C96-TNB, the fluorescence lifetime distributions obtained at the different GdnHCl concentrations, were bimodal (Figure 4). Lifetimes between 0.001 and 0.6 ns, were taken to arise from native-like (N-like) forms, and lifetimes > 0.6 ns, were taken to arise from unfolded-like (U-like) forms of the protein. This assignment was based on the observation that the U state predominantly populates lifetimes > 0.6 ns (U-like) and the N state predominantly populates lifetimes < 0.6 ns (N-like).^12–13^ Such grouping of lifetimes allowed the fractions of molecules in the U-like and N-like sub-populations to be determined at each GdnHCl concentration (see Materials and Methods). For each protein variant, the dependence of the MEM-derived fraction of molecules that were U-like (see Materials and Methods) was found to match the unfolding transition monitored by the measurement of fluorescence intensity or mean fluorescence lifetime (Figures 2, 5 and S3) and yielded similar values for the stability, ΔG_U_ and the midpoint of the transition, C_m_ (Figure 5 and Table S1). This was important because it validated the grouping into N-like and U-like sub-populations.

**Figure 4.**
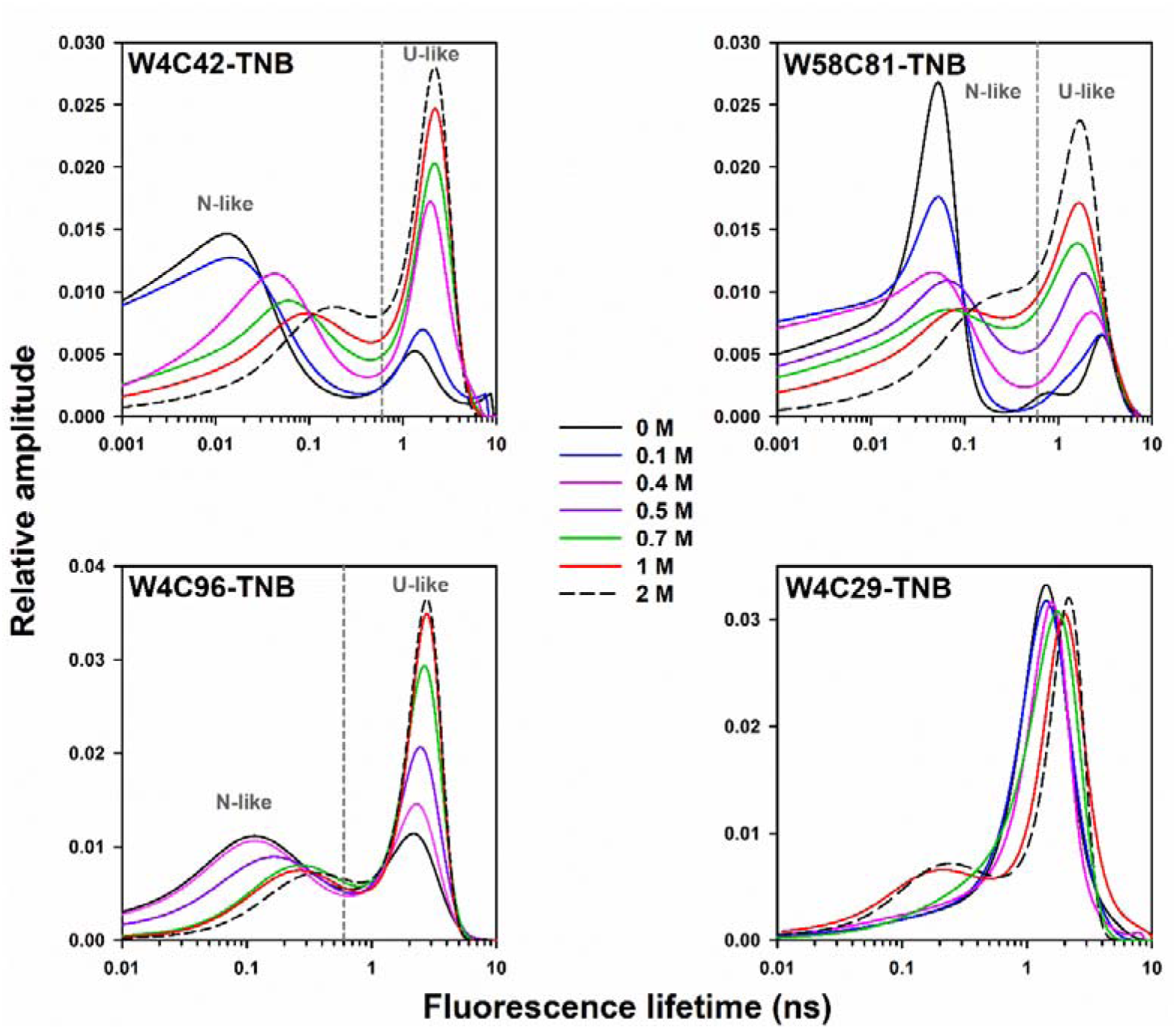
MEM-derived fluorescence lifetime distributions of the TNB-labeled variants of dcMN at different GdnHCl concentrations. The differently colored lines correspond to various concentrations of GdnHCl, as indicated.

**Figure 5.**
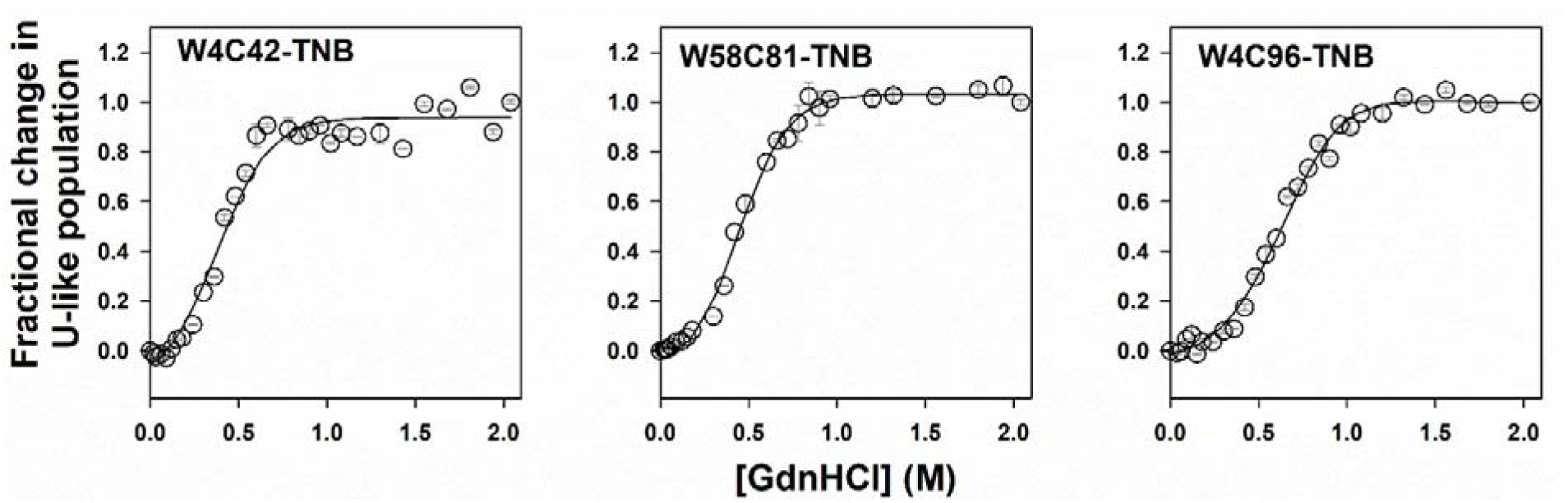
Fractional change in the U-like population calculated from the relative sum of amplitudes of the MEM distributions at different concentrations of GdnHCl. The solid line passing through each dataset is a non-linear, least-squares fit to a two-state unfolding model.^41^ The fits gave values for ΔG_U_ of 9.3 ± 0.2, 9.4 ± 0.3 and 10.6 ± 0.1 kcal mol^-1^ for W4C42-TNB, W58C81-TNB and W4C96-TNB, respectively. The mid-points (C_m_) of the unfolding transitions for W4C42-TNB, W58C81-TNB and W4C96-TNB are at 0.28 ± 0.04, 0.33 ± 0.06 and 0.61 ± 0.02 M, respectively. The error bars represent the spread in the data obtained from two independent experiments.

The main features of the fluorescence lifetime distributions were as follows: (a) for the N-like forms, the peak position shifted gradually towards longer lifetimes with increasing GdnHCl concentration; (b) for the U-like forms, the peak shifted towards longer lifetimes in the case of W4C42-TNB, and towards shorter lifetimes in the case of W58C81-TNB; (c) the magnitude of these shifts varied among the different TNB-labeled variants (Figures 4 and S4); and (d) the relative fractions of N-like and U-like sub-populations changed in an apparently two-state manner (Figure 5), with an increase in the U-like fraction and a corresponding decrease in the N-like fraction as GdnHCl concentration increased. Thus, both gradual and all-or-none changes were observed within chain B, and chain A, as well as across the two chains.

In contrast, the lifetime distributions for the helix segment (W4C29-TNB) remained unimodal throughout the transition (Figure 4). The peak position shifted gradually towards longer lifetimes with increasing GdnHCl concentration, indicating a gradual change in the intra-helix distance during the unfolding transition (Figures 4 and S4). It should be noted that for all the unlabeled proteins, the fluorescence lifetime distributions remained unimodal throughout the unfolding transition (Figure S5) with the peaks not showing any significant shift upon unfolding (insets, Figure S5).

The dependence of the MEM peak lifetime on GdnHCl concentration was clearly sigmoidal for the N-like subensembles seen for W4C42-TNB, W58C81-TNB and W4C96-TNB (Figure S4). In contrast, the MEM peak lifetime for the U-like subensembles seen for these variants, as well as for W4C29-TNB, showed non-sigmoidal dependences on GdnHCl concentration (Figure S4), pointing to non-cooperativity. In order to interpret these changes in the MEM peak lifetimes of the TNB-labeled variants, the distances within chain B (in W4C42) and chain A (in W58C81), across both chains (in W4C96), and spanning the helix in chain B (in W4C29) were calculated using equation 7 (Materials and Methods) after correcting for the small peak shifts of the unlabeled proteins (Figures 6, S4 and S5).

**Figure 6.**
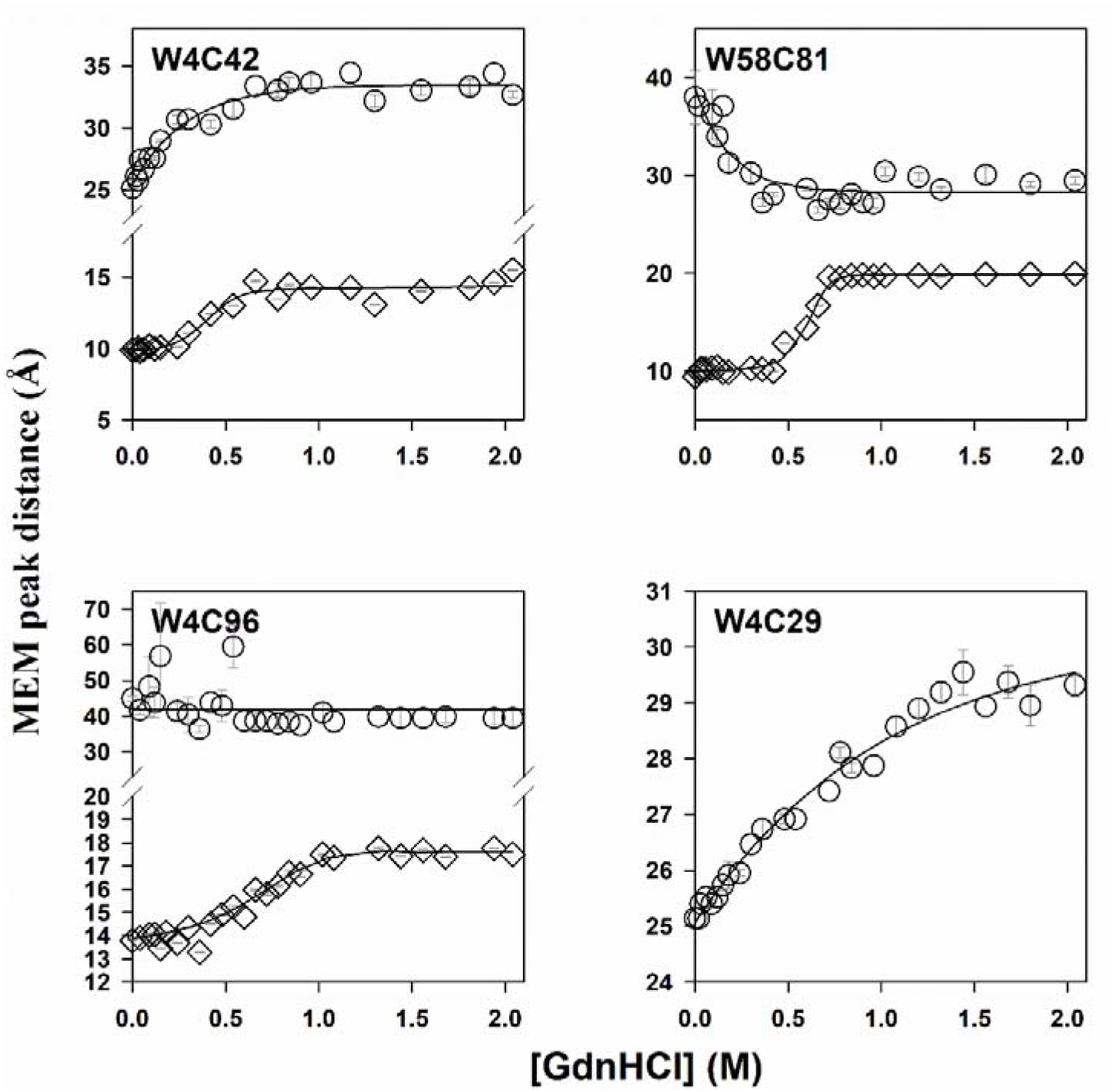
Change in the dimensions of the N-like and U-like sub-populations with increasing GdnHCl concentration. The MEM peak lifetimes for the TNB-labeled proteins and for the corresponding unlabeled proteins were used to calculate the MEM peak FRET efficiencies (equation 6, see Materials and Methods), which were converted to distances (equation 7, see Materials and Methods). Diamonds and circles represent the distances corresponding to the peaks of the MEM distributions of the N-like and U-like sub-populations, respectively. The error bars represent the spread in the data obtained from two independent experiments.

While the MEM peak lifetime of the U-like sub-population observed using the FRET pair mapping separation across the two chains (Trp4 -C96TNB) shifted slightly toward longer lifetimes (Figure 4), a comparable shift was also observed for the fluorescence lifetime distributions of the corresponding unlabeled protein (Figure S5). Consequently, the U-like sub-population exhibited negligible FRET with no discernible change over the entire range of GdnHCl concentrations. The intra-molecular distances inferred from the very small FRET values exceeded ∼40 Å, at all GdnHCl concentrations, indicating that the chains had physically separated even in zero denaturant. It should be noted that the separation of Trp4 from Cys96-TNB in MNEI at high GdnHCl concentration is less.^12^

The negligible FRET observed in the U-like sub-population for dcMN was consistent with the ensemble-averaged FRET efficiency being near-zero at GdnHCl concentrations > 1.5 M (Figures 2 and 3). Only at low GdnHCl concentrations, at which the U-like sub-population was negligibly populated (Figure 5), did the ensemble-averaged FRET efficiencies have significant values (Figures 2 and 3) because they were dominated by the contribution of the N-like sub-population. Hence, the ensemble-averaged FRET analysis was consistent with the MEM analysis which suggested that the two chains have separated even in the absence of GdnHCl in the U-like subpopulation.

To assess the extent of deviation of the four protein variants from two-state unfolding behaviour, the MEM distribution at each GdnHCl concentration was fitted to the weighted sum of the N- and U-state fluorescence lifetime distributions (Figure S6) using equation 5 (see Materials and Methods). In the case of W4C42-TNB, in which a distance change within chain B was monitored, and W58C81-TNB, in which a distance change within chain A was monitored, it was found that fluorescence lifetime distributions in the transition region did not fit to the weighted sum of the N state and U state fluorescence lifetime distributions (Figures S6 and S7). Deviations were also observed in the case of W4C96-TNB, in which a distance across the two chains was monitored and W4C29-TNB, in which a distance spanning the helix of chain B was monitored, but they were of much smaller magnitude (Figure S7). It should also be noted that none of the protein variants showed an iso-lifetime point, that is, a single fluorescence lifetime at which all distributions intersected. The absence of such a point, expected for a two-state transition, further confirmed that the unfolding of dcMN is not strictly two-state, but involves heterogeneous ensembles populated throughout the transition.

### Motional dynamics of Trp4 (in W4C42) and Trp58 (in W58C81) revealed heterogenous unfolding

The unfolding transition monitored by steady-state anisotropy (r_ss_) was found to be identical to that monitored by fluorescence intensity (Figure S8b). Values of r_ss_ were also obtained from the time-resolved anisotropy decays at different GdnHCl concentrations (Figure S9) and were found to coincide with the steady-state fluorescence anisotropy measured directly (Figure S8b), supporting the accuracy of the time-resolved measurements.

Since r_ss_ is a function of both the rotational dynamics of the fluorophore and its excited-state lifetime (equation 12, Materials and Methods), a change in it (Figure S8b) may not reflect a change in rotational dynamics alone. Hence, the different rotational correlation times obtained from time-resolved fluorescence anisotropy decays of Trp4 in W4C42 and of Trp58 in W58C81 (Figure S9) were analyzed to obtain direct information on the rotational dynamics, which in turn relate to the structural compactness around Trp4 and Trp58 in chains B and A, respectively (Figure 7). Trp58 in chain A also forms a part of the core β2-β3 dimer interface (Figure S8a), and thus is expected to offer insights into the structural and dynamical aspects not only within chain A but at the β2-β3 dimeric interface as well.

**Figure 7.**
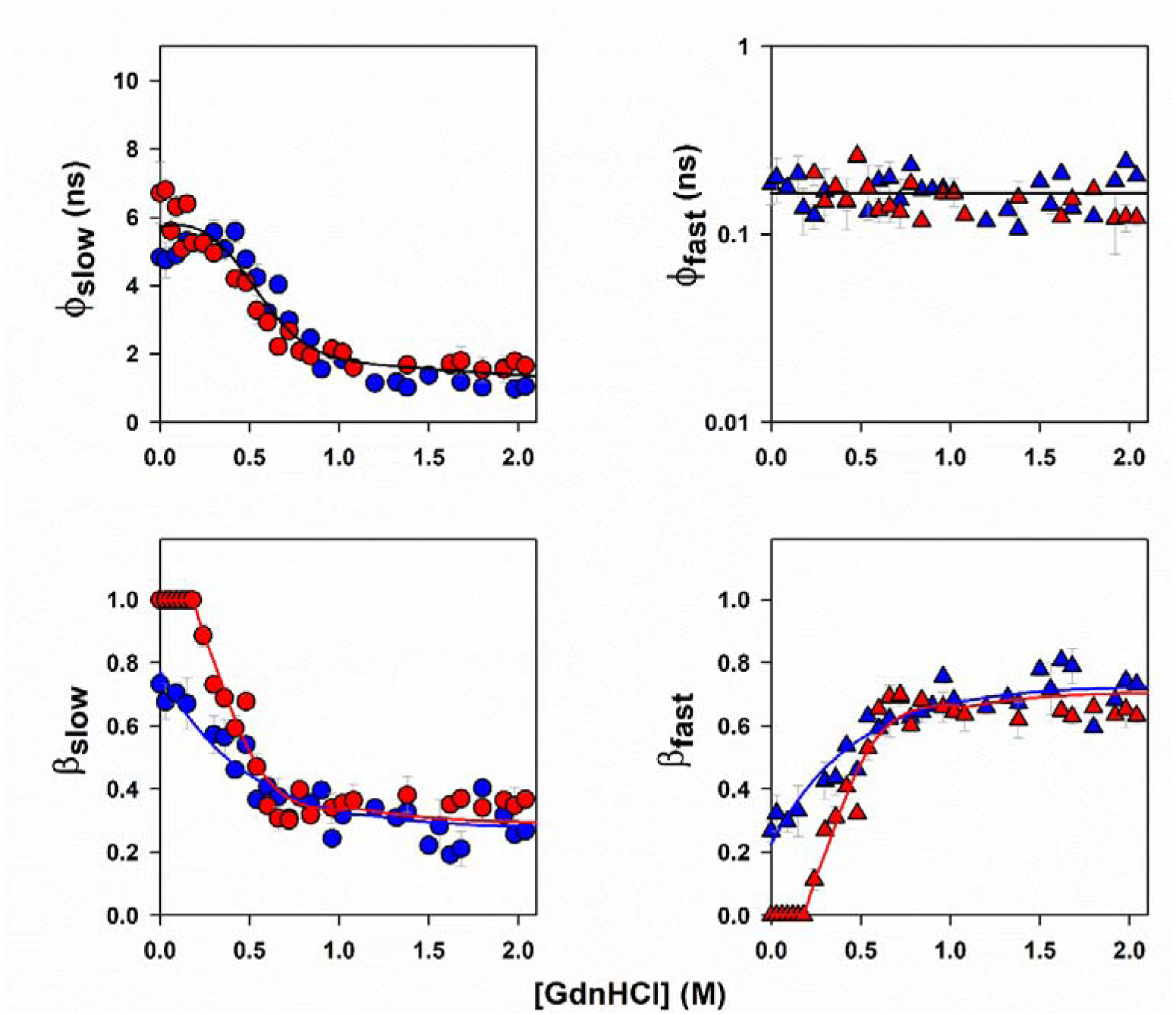
Equilibrium unfolding of W4C42 and W58C81 monitored by time-resolved anisotropy measurement. Data for Trp4 (blue) were obtained by measurements on W4C42, and data for Trp58 (red) were obtained from measurements on W58C81. Dependence of the slow (circles) and fast (triangles) rotational correlation times, along with their corresponding relative amplitudes, on GdnHCl concentration is shown. The error bars represent the spread in the data obtained from two independent experiments.

Fluorescence anisotropy can decay relatively slowly by global rotational diffusion of the protein and/or rotational diffusion of a protein segment, defined by a slow rotational correlation time, □_slow_ with corresponding relative amplitude β_slow_ as well as by independent fast rotational motion of the Trp side-chain, defined by a fast rotational correlation time, □_fast_ with corresponding relative amplitude β_fast_. The site-specific time-resolved fluorescence anisotropy decay measurements revealed distinct local dynamic environments for the two Trp residues in the native protein. In 0 M GdnHCl, the anisotropy decay curve of Trp4 (in W4C42) required a two-exponential fit, yielding □_slow_ = ∼ 6 ns, and □_fast_ = ∼ 0.2 ns, with β_fast_ = 0.25 (Figures 7 and S9). In contrast, the anisotropy decay curve of Trp58 (in W58C81) fit well to a single-exponential equation, yielding □_slow_ (Figure 7). For both Trp residues, the value of □_slow_ (6 ± 1 ns) was that expected for an ∼ 11 kDa protein in aqueous media at 25°C.^50–51^ The presence of a measurable fast component for Trp4 indicated that it undergoes local side-chain motion in the N state, which Trp58, located at the dimer interface (Figure S8a), does not. This indicated that Trp58 is in a conformationally restrictive environment with its side-chain immobilized in the native structure. It should be noted that the values of initial fluorescence anisotropy values (r□) remained constant across the entire GdnHCl range for both Trp residues (Figure S10). This indicated that the orientations of the absorption and emission transition dipole moments did not change appreciably during unfolding, and, consequently, variations in the fluorescence anisotropy decays arose from changes in rotational dynamics rather than changes in photo-selection or fluorophore geometry.

The value of □_slow_ was observed to decrease in a sigmoidal manner with increasing GdnHCl concentration for both Trp4 (in W4C42) and Trp58 (in W58C81). This decrease reflects a transition from rigid-body global tumbling to dynamics increasingly dominated by local segmental motions of the Trp-containing segments. In the U state, fluorescence anisotropy of both Trp residues decayed by segmental motion with a rotational correlation time of 1 ns (Figure 7), as well as the by fast rotational motions of the side-chains. The overlap in the cooperative sigmoidal transitions mapped by the values of □_slow_ for Trp4 and Trp58, suggested simultaneous global loosening of the two chains.

The value of □_fast_for Trp4 remained invariant at about 0.2 ns across the entire range of GdnHCl concentration. For Trp58, □_fast_ could be resolved only at GdnHCl concentrations above 0.2 M, and it too remained invariant at 0.2 ns (Figure 7). In the case of both Trp residues, β_fast_ was found to increase progressively in an apparently exponential manner with increasing denaturant concentration, consistent with a gradual increase in local motional freedom, before reaching a limiting value of about 0.65. This continuous redistribution between global tumbling and local flexibility for both Trp4 and Trp58 suggests a heterogeneous and distributed structural loosening, rather than an all-or-none transition. Together, these observations demonstrated that the unfolding process cannot be described by a simple two-state model but instead involves intermediate motional modes and progressive structural disruption.

## Discussion

### Equilibrium unfolding of dcMN is not cooperative

Fluorescence intensity measurements of the equilibrium unfolding of all four unlabeled protein variants and their labeled counterparts, as well as steady-state FRET measurements mapping expansion of chain B (in W4C42), chain A (in W58C81), and the inter-chain distance (in W4C96), would suggest that the GdnHCl-induced unfolding of dcMN occurs in a cooperative manner (insets, Figure 2). Nevertheless, the unfolding of the segment mapping the helix (W4C29) appears to be gradual. Indeed, the lack of cooperativity in the unfolding transition become more evident in tr-FRET measurements upon a population level MEM analysis. Considerable heterogeneity is revealed. Unfolding is seen to proceed through sub-populations of N-like forms expanding continuously while transiting into U-like forms which also expand with an increase in GdnHCl concentration (Figure 6). The observation that the MEM-derived distributions of fluorescence lifetimes for all the protein variants cannot be described adequately as the weighted sum of the N and U state distributions (Figures S6 and S7), provides strong evidence that the cores of both chains, the inter-chain interface and the helix, all unfold in a non-cooperative manner.

### Different regions of the protein display significant differences in cooperativity

The observation that the segment spanning the helix exists in a single conformational ensemble which expands continuously with an increase in GdnHCl concentration (Figure 6) suggests that the unfolding of the helix occurs in a completely gradual manner. In contrast, the observation of co-existing sub-populations of N-like and U-like forms, which differ in their mean intra-chain and inter-chain dimensions, for segments mapping the cores of chain A (in W58C81) and chain B (in W4C42) and across the two chains (in W4C96), suggests that they are separated by a significant free-energy barrier. A previous HX-MS study had shown that at equilibrium, multiple partially unfolded intermediate ensembles, separated by significant energy barriers, coexist with the N and U states. Moreover the structures differed in the absence and in the presence of GdnHCl.^33^ For example, the most U-like intermediate ensemble had only β2 structured in zero denaturant, and both β2 and β3 structured in the presence of low concentrations of GdnHCl, but the HX-MS studies could not determine whether this difference arose from local or non-local interactions. In the case of adenylate kinase too, the distribution of cooperatively exchanging intermediates could be modulated by denaturant,^52^ and in the case of barstar^8^ and the SH3 domain of PI3K,^30^ β-sheet regions were found to unfold relatively more cooperatively than other structural elements under equilibrium conditions.

Nevertheless, in the case of dcMN, while different segments mapping inter-β strand distances transit from a N-like to a U-like sub-population in an apparently cooperative manner (Figure 5), distances within each sub-population are seen to change in an apparently continuous manner with a change in GdnHCl concentration (Figure 6). In the previous HX-MS study too, it was seen that each intermediate ensemble consisted of molecules sampling many different conformations that differed as little as in having or not having structure at only one amide site.^33^

### Swelling of the partially contracted N-like sub-populations is cooperative

The observation that the N-like sub-populations undergo swelling in an apparently cooperative manner with increasing GdnHCl concentration, both for the intra-chain segments in chains A (in W58C81) and B (in W4C42) and for the segment spanning both chains (inW4C96) (Figure 6), suggests that molecules contracted at these segments respond in a coordinated manner to the disruption of stabilizing interactions. Importantly, the denaturant dependence of swelling of the N-like ensemble differs across sites: the expansion transitions for W4C42 and W58C81 saturate at ∼ 0.7 M GdnHCl, whereas the corresponding transition for W4C96 is broader and saturates only at ∼1 M GdnHCl (Figure 6). This difference suggests that W4C42 and W58C81 report predominantly on the sub-global stabilities of the N-like sub-populations monitored in chains B and A, respectively, which are determined primarily by less stable intra-chain contacts that are disrupted at lower denaturant concentrations. On the other hand, W4C96, in which the FRET-monitored distance spans both chains, reports on the global stability of the N-like sub-population of dcMN, which is determined not only by intra-chain contacts but also by stabilizing interactions that couple the two chains together. It appears that the N-like forms preserve an inter-dependent network of non-local contacts, including β-sheet hydrogen-bonding networks that link together distant sequence positions^53^ as well as stabilizing non-covalent interactions at the inter-chain interface. Similar coupling between non-local contacts and cooperative expansion has been inferred in other β-rich proteins, where high contact order and β-sheet topology impose free-energy barriers that must be crossed collectively.^54–57^ Native-state HX-NMR studies have shown that groups of residues can lose protection in concert, defining cooperative foldon units,^58^ and a kinetic HX-MS on the SH3 domain of PI3 kinase reveal that hydrogen bonds between adjacent strands rupture collectively rather than strand by strand.^59^

### The U-like sub-populations undergo non-cooperative unfolding

The observation that in the case of the inter-chain distance segment (W4C96), negligible FRET is observed for the U-like sub-population across the entire range of GdnHCl concentrations, suggests that the two chains have separated even at 0 M GdnHCl (see Results). Consequently, the changes in the dimensions of individual chains A (monitored in W58C81) and B (monitored in W4C42) occur in a continuous, non-cooperative manner, with increasing denaturant concentration.

The continuous expansion of the segment mapping the core of chain B (in W4C42) with increasing GdnHCl concentration is consistent with behavior expected when solvent–chain interactions dominate over intra-chain stabilizing interactions. Such monotonic expansion has been observed for many intrinsically disordered proteins,^60–61^ which tend to swell with increasing denaturant concentration as solvent quality improves and chain–solvent interactions increasingly outweigh chain–chain interactions. Similar expansion of the U state ensemble with increasing denaturant concentration has been reported for a wide range of globular proteins under unfolding conditions.^8, 62–63^

In contrast, the behavior of chain A (monitored in W58C81) is markedly different. Surprisingly, the segment mapping the core of chain A is seen to undergo gradual contraction upon addition of GdnHCl. In 0 M GdnHCl, this segment has a size of ∼38 Å and is 35% expanded relative to its dimension in the U state at high GdnHCl concentration. This would suggest that water is a good solvent for unfolded chain A, as has been suggested for the unfolded forms of globular proteins.^64^ However, a solvent quality based interpretation is insufficient to account for this behaviour.

This observation can be rationalized by electrostatic repulsion playing a dominant role in determining the dimensions of the U-like forms of chain A at low denaturant concentrations. Screening of electrostatic interactions by Gdn□ and Cl□ ions would reduce repulsive interactions between like-charged residues, allowing weak hydrophobic and non-local interactions to promote compaction. Similar salt-induced contraction of unfolded protein has been reported previously, including in intrinsically disordered proteins, where increasing ionic strength leads to chain compaction through electrostatic screening.^65–66^ Low concentrations of GdnHCl have also been shown to induce compaction of the denatured states of apomyoglobin and cytochrome c.^67^ Furthermore, perturbation of even a single electrostatic interaction by mutation has been shown to significantly alter U state dimensions in the absence of any denaturant, and screening of charge by the addition of 0.5 M salt concentration was found to restore the U-state dimensions to that of the wild-type U state ensemble.^68^

It has been suggested that the net charge per residue (NCPR), defined as the net charge normalized by sequence length, is a key determinant of the global dimensions of unfolded and intrinsically disordered polypeptide chains.^69–70^ Both chains A and B fall within the weak polyampholyte (Janus sequence) boundary region in the diagram of states determined by the CIDER (Classification of intrinsically disordered ensemble relationships) program.^69–70^ In this regime, electrostatic interactions are context-dependent and do not enforce a single U state conformation but can stabilize either expanded or compact ensembles. In this regime, small imbalances between repulsive and attractive electrostatic interactions are sufficient to bias the unfolded ensemble toward expansion or compaction, as established by the classical polymer theory of polyampholytes and by studies on intrinsically disordered protein sequences.^70–71^ The sequence segment mapping the core of chain A exhibits a small but finite net charge bias (NCPR ≈ -0.08; net charge ≈ -2 at pH 8), resulting in a weak bias toward repulsive electrostatic interactions and a more expanded ensemble in 0 M GdnHCl. In contrast, the sequence segment mapping the core of chain B is charge-neutral (NCPR ≈ 0; net charge ≈ 0 at pH 8), a condition under which polyampholytes can adopt compact conformations.^71–72^ Screening of electrostatic interactions upon addition of GdnHCl therefore leads to contraction of chain A and expansion of chain B in the low denaturant regime.

These observations indicate that the inter-chain interface plays a central role in modulating unfolding cooperativity in dcMN by restricting chain entropy. When the interface is intact (as in the N-like sub-population; see above), it promotes coordinated, cooperative responses across both chains. In contrast, in the U-like sub-population, where the inter-chain interface is already disrupted, release of this entropic constraint abolishes coupling, allowing the two chains to undergo chain-specific, non-cooperative structural rearrangements. Thus, this study demonstrates directly that unfolding cooperativity is organized at the level of intermediate sub-populations and is regulated by connectivity-imposed chain entropy. Previous studies on coiled-coil heterodimers^73^ and PDZ–ligand complexes^74^ had shown that interfacial contacts can drive coordinated, all-or-none unfolding of coupled regions, whereas more weakly coupled segments display non-cooperative behaviour,^75–76^ but could not resolve how the degree of cooperativity is selected for within heterogeneous intermediate ensembles.

### Time-resolved fluorescence anisotropy decay measurements reveal site-dependent and asynchronous changes in local motions during unfolding

The relative amplitudes of the fast and slow decay components (β_fast_ and β_slow_) report the fraction of molecules that lose anisotropy *via* fast local Trp motion *versus* slower segmental or global motions (Figure 7). In a strictly two-state unfolding process, local probes located in different regions of the protein would therefore be expected to exhibit similar denaturant dependences of these relative amplitudes. Instead, Trp4 and Trp58 show markedly different behaviors. For Trp4, the relative contribution of fast local motion (β_fast_) increases gradually and in an approximately exponential manner with increasing GdnHCl concentration, indicating a continuous redistribution of the ensemble toward conformations with increasing local flexibility. In contrast, for Trp58, β_fast_ remains zero at low denaturant concentrations and increases gradually only beyond ∼ 0.2 M GdnHCl, suggesting that local motional freedom at this site does not occur in the N state. These differences demonstrate that distinct regions of dcMN lose motional constraints at different denaturant concentrations, revealing heterogeneous and asynchronous structural loosening rather than a concerted all-or-none transition.

Moreover, if both fast and slow depolarization pathways were accessible simultaneously to all molecules, the relative amplitudes would be dictated solely by the corresponding rotational correlation times. For example, in the U state, where □_slow_is 1 ns and □_fast_is 0.2 ns, the value of β_fast_ expected for such a system would be approximately 0.83 (1/1.2). Instead, the observed value of β_fast_ is ∼0.65. This discrepancy indicates that in the U state, and similarly in the N and intermediate states, there exist minor sub-populations (∼15% in the U state) in which the Trp side-chain remains dynamically constrained and cannot undergo independent local motion. Thus, the anisotropy data establish that unfolding proceeds through a progressive redistribution among sub-populations with different local motional constraints and cannot be described by a simple cooperative two-state model. Similar deviations from simple two-state behavior have been revealed by time-resolved fluorescence anisotropy decay measurements of barstar, for which the rotational dynamics in the unfolding transition region could not be described as a weighted sum of the native and unfolded state decays.^77^ In the case of α-subunit of tryptophan synthase, non-monotonic changes in rotational correlation times were observed during folding, indicating asynchronous rather than cooperative structural rearrangements.^78^ Similarly, studies on tubulin,^79^ creatine kinase,^80^ and yeast glutathione reductase^81^ identified intermediate species and gradual structural loss, consistent with multi-step rather than cooperative unfolding.

### Comparison of the unfolding cooperativity of dcMN and MNEI

Although dcMN and MNEI have nearly identical native structures (Figure 1a), their unfolding behavior differs markedly when resolved at the level of MEM-derived sub-populations (Figure S11). This isolates how covalent chain connectivity, via restriction of chain entropy, modulates unfolding cooperativity independent of native structure.

For the helix segment (monitored in W4C29), the unfolding transition is gradual for both MNEI and dcMN, indicating that the non-cooperative unfolding of the helix is intrinsic to the helix itself and largely insensitive to differences in chain connectivity or overall protein topology. In contrast, connectivity-dependent differences are observed for the β-sheet core (monitored in W4C42) and inter-chain segments (monitored inW4C96/W4C97). The N-like sub-populations swell via similar cooperative transitions, for both MNEI and dcMN. In contrast, it is seen that the U-like sub-population observed for W4C42 unfolds non-cooperatively in the case of dcMN but cooperatively in the case of MNEI. For the inter-chain segment monitored in W4C96: unfolding of the U-like sub-population is cooperative in the case of MNEI, but the U-like sub-population is as expanded as the U state in the case of dcMN.

Overall, these comparisons show that in the case of MNEI, unfolding transitions remain largely coupled at the segmental level, consistent with a covalently continuous polypeptide in which interacting structural elements remain effectively tethered even upon partial expansion. In sharp contrast, dcMN exhibits a clear separation of cooperative and non-cooperative behavior between sub-populations. Covalent continuity in the MNEI chain maintains cooperative unfolding across both the N-like and U-like sub-populations, whereas in the case of dcMN, cooperativity is selectively preserved only within the N-like sub-population and lost within the U-like ensemble, which unfolds in a chain-specific, non-cooperative manner. Thus, covalent linkage between interacting structural elements stabilizes cooperative responses even within partially expanded ensembles, whereas its absence permits uncoupled, non-cooperative behavior.

This contrasting behavior can be rationalized in terms of chain connectivity and effective concentration. The β2–β3 interface is structurally equivalent in the native structures of dcMN and MNEI; however, in dcMN, the interacting β strands reside on separate polypeptide chains, whereas in MNEI they are covalently linked within a single chain (Figure 1a). In the case of dcMN, the effective concentrations governing various inter-chain interactions will be low because the partners are on different chains, and the high entropic cost of bringing the interacting partners together lowers the strengths of the stabilizing interactions. Consequently, unfolding is less cooperative. In the case of MNEI, effective concentrations governing the same interactions are higher because the partners are on one chain. Consequently, the stabilizing interactions are stronger and loss of structure during unfolding is cooperative. In the case of circularly permuted proteins too, the changes in covalent connectivity alone can modulate (un)folding cooperativity, even when native structure is preserved.^82–85^ It therefore seems likely that the lack of cooperativity seen for the U-like sub-populations of dcMN arises from the two-chain topology and the loss of covalent linkage between interacting β-strand elements. Given the prevalence of coupled structural elements in multidomain and multimeric proteins, this mechanism is likely to be broadly relevant for regulating unfolding cooperativity in complex protein systems.

## Conclusion

In summary, the present study has identified chain connectivity as a molecular determinant of unfolding cooperativity in monellin. Despite possessing nearly identical native structures, the double-chain and single-chain variants exhibited distinct unfolding behavior when resolved at the level of intermediate sub-populations. In dcMN, effective inter-chain coupling preserved cooperative expansion within N-like sub-populations, whereas the loss of this coupling in U-like sub-populations promoted pronounced chain-specific, non-cooperative behavior during unfolding. In contrast, covalent continuity in MNEI enforces cooperative unfolding across both the N-like and U-like sub-populations by restricting chain entropy. Collectively, these findings demonstrate that unfolding cooperativity is governed by connectivity-imposed coupling between structural elements and can be selectively lost within specific intermediate sub-populations rather than uniformly throughout the unfolding process.

## Materials and Methods

### Protein expression, purification, and TNB labelling

The wild-type protein (W4C42) has a single tryptophan (Trp4) residue located in the first β-strand and a single buried cysteine (Cys42) in the second β-strand of chain B. Site-directed mutagenesis was used to generate the single-tryptophan, single-cysteine containing mutant variants: C42CAQ29C (W4C29), C42AP96C (W4C96), and W4YY58WC42AT81C (W58C81). Protein purification was carried out following an established protocol.^40^ The single cysteine residue in each protein variant was conjugated with the acceptor thionitrobenzoate (TNB) moiety. Labeling was performed as previously described.^21^ In brief, the protein was unfolded in 2 M GdnHCl and incubated with DTNB solution at pH 8 for ≥ 2 h; after which excess label was removed by desalting. The purity of each protein preparation was confirmed by electrospray ionization mass spectrometry (ESI-MS). The mass of the labeled protein showed the expected increase of 197 Da, corresponding to the mass of the TNB group, and the extent of labeling was > 95%. The concentrations of the unlabeled protein solutions were determined from absorbance measurements at 280 nm, using the molar extinction coefficient value of 14600 M^-1^ cm^-1^. For the TNB-labeled proteins, the contribution of the TNB absorbance to the total absorbance at 280 nm was 20% for all TNB-labeled protein variants, except for W4C96-TNB, for which the contribution was 10%.^12^

### Reagents

All the experiments were carried out at pH 8.0 and 25°C. The reagents used in the experiments were of the highest purity grade from Sigma. Guanidine hydrochloride (GdnHCl) was purchased from Thermo Scientific, and was of the highest purity grade. The native buffer contained 50 mM Tris and 250 µM EDTA, and unfolding buffer contained, in addition, GdnHCl. 1 mM DTT was added for all experiments with unlabeled proteins. The concentrations of GdnHCl solutions were determined by the measurements of the refractive index on an Abbe 3L refractometer from Milton Roy. All buffers and solutions were passed through 0.22 µm filters before use. A protein concentration of 5 µM was used for all experiments.

### Measurement of fluorescence and far UV CD spectra

Fluorescence spectra were collected on a Fluoromax 4 (Horiba) spectrofluorometer. The protein samples were excited at 295 nm, and the emission spectra were collected from 310 to 450 nm, with a response time of 1 s, and excitation and emission bandwidths of 1 nm and 5 nm, respectively. Each spectrum was recorded as the average of three fluorescence emission wavelength scans. Measurements of CD spectra were carried out on a Jasco J-815 spectropolarimeter. Far-UV CD spectra were collected using a 0.2 mm pathlength cuvette, with a bandwidth of 1 nm, a response time of 1 s, and a scan speed of 20 nm/min. Thirty wavelength scans were averaged for each spectrum.

### Steady-state fluorescence and anisotropy measurements

For equilibrium unfolding studies, protein samples were incubated in different concentrations of GdnHCl (0 to 2 M) for ≥ 48 h prior to measurement. The fluorescence intensities of both unlabeled and TNB-labeled protein variants for steady-state FRET measurements were measured using the MOS-450 optical system (Biologic). The protein samples were excited at 295 nm, and the emitted fluorescence was monitored at 360 nm using a 10 nm band-pass filter (Asahi Spectra). The excitation slit width was kept at 2 nm. For each sample, the data were acquired and averaged for 30 s. The fluorescence anisotropies of Trp4 in W4C42 and Trp58 in W58C81 were measured using a Fluoromax 4 (Horiba) spectrofluorometer by monitoring the emission at parallel and perpendicular polarizations simultaneously using the T-format optical arrangement.

### Time-resolved fluorescence and anisotropy measurements

Fluorescence lifetime decay curves for the four pairs of unlabeled and labeled proteins, and fluorescence anisotropy decay curves for unlabeled W4C42 and W58C81 were acquired after equilibrating the samples at different GdnHCl concentrations for ≥ 48 h prior to measurements.

Fluorescence lifetime and anisotropy measurements were carried out using an excitation wavelength of 295 nm, generated by frequency-tripling 885 nm femtosecond pulses from a Ti:sapphire laser. Emission at 360 nm was detected using hybrid PMTs (HPM-100-40 or HPM-100-07). Time-correlated single-photon counting (TCSPC) was employed to record decay profiles, and instrument response functions (IRF) were recorded for deconvolution. A polarizer (Glan-Thompson), set at 54.71° (magic angle) with respect to excitation, was kept in front of the long-pass filter to avoid polarization effects. In time-resolved anisotropy decay measurements, the emission was collected at directions parallel (I_‖_) and perpendicular (I_┴_) to the polarization of the excitation beam, for the same duration. The IRF was measured using light scattered from a LUDOX solution. The HPM-100-07 consistently gave an IRF FWHM of ∼60 ps, and the HPM-100-40 detector gave an IRF FWHM of ∼100 ps. The fluorescence lifetime of NATA (2.5–2.8 ns) in water was routinely recorded as a standard. All decay curves were collected to a peak count of 20 000, and up to 99.9% of completion. Detailed descriptions of the instrumentation for lifetime^12^ and anisotropy^77, 86–87^ measurements have been reported previously.

### Data analysis

#### Analysis of the fluorescence lifetime decay traces

Details of the analysis are given in an earlier study.^12^ A brief description is given below:

#### Discrete analysis

The decay traces were fit to a sum of exponentials,

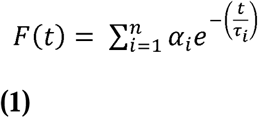

Here, α_i_ is the relative amplitude of the τ_i_ lifetime component, t is time, and n ranges from 3 to 4.

The amplitude-weighted mean lifetime, τ_m_ was caculated as:

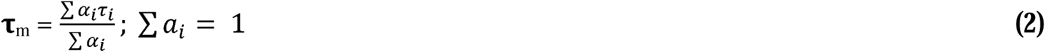

#### MEM analysis

The analysis began with the assumption that the decay corresponds to a distribution of 100-150 lifetimes in the range 10 ps to 10 ns. The *a priori* distribution of lifetimes was assumed to be uniform in the logarithms of lifetimes being uniformly distributed in this range. Then, the best fit values of α_i_and τ_i_ were determined using the Maximum Entropy Method (MEM).^13, 21^

The *a posteriori* distribution of these parameters was obtained by maximizing the Shannon Jaynes entropy S, defined as

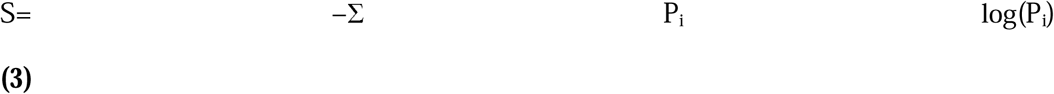

P_i_ is the normalized amplitude of the ith lifetime.

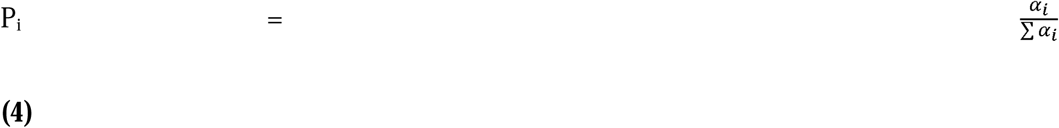

α_i_ is the amplitude of the ith lifetime.

Analysis parameters were optimized for obtaining precise and reproducible MEM distributions. The most probable (MEM peak) lifetime refers to the lifetime corresponding to the maximum amplitude in the lifetime distribution data.

### Fitting of MEM analysis-derived distributions to a two-state model

Fluorescence lifetime distributions obtained from MEM analysis were fitted to a two-state model using a nonlinear least-squares curve fitting algorithm implemented in MATLAB. This algorithm used the fluorescence lifetime distributions of the native state, N(⍰), and the unfolded state, U(⍰), as basis distributions, and carried out a nonlinear least-squares fit across all GdnHCl concentrations according to the equation:

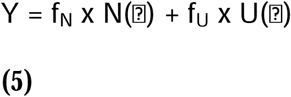

where f_N_ and f_U_ represent the fractions of native and unfolded protein, respectively, at each GdnHCl concentration. The root mean-squared deviation (rmsd) at a given GdnHCl concentration (Figure S7) was calculated as the square root of the mean of the squared residuals across all lifetime values.

### FRET efficiency and distance determination

The mean FRET efficiency (<E_FRET_>) was obtained from mean fluorescence lifetimes of the unlabeled (<τ_D_>) and the corresponding TNB-labeled (<τ_DA_>) variants using the following equation:

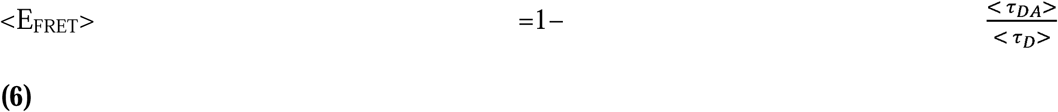

The mean fluorescence lifetimes were determined by discrete analysis of the fluorescence decay traces as described earlier.^12, 21^

The mean FRET efficiency value was converted to the average intramolecular distance (<R_DA_>) using the F□rster equation:

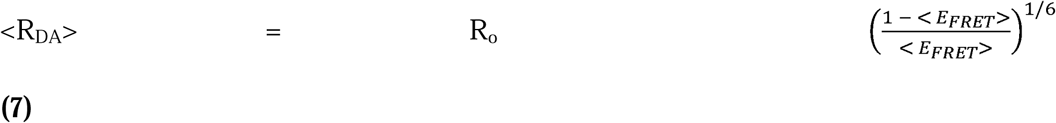

The values for the Forster’s distance, R_o_ used for the different protein variants are listed in Table S2 and were determined as described previously.^12^

### Determination of the fractions of molecules in the N-like and U-like subpopulations from MEM distributions

To determine the fractions of N-like and U-like molecules from the MEM distributions (see introduction) at different GdnHCl concentrations, it was essential to take into account the observation that, for any segment, the equilibrium U state contained a fraction of molecules exhibiting lifetimes corresponding to an N-like distribution (<0.6 ns), while the equilibrium N state included a fraction of molecules exhibiting lifetimes corresponding to a U-like distribution (>0.6 ns) (Figures 4 and S5). These fractions were comparable for each pair of unlabeled (Figure S5) and corresponding labeled (Figure 4) unfolded proteins, suggesting that they originate from differences in the electronic structure of the fluorophore or distinct Trp rotamers^88–91^ rather than from the presence of the quenching TNB moiety. The fraction of molecules expanded (U-like) at a given segment was determined using the equation: f_U_=Y_i_−Y_N_/Y_U_−Y_N_, where Y_U_ represents the relative sum of amplitudes for the U-like distribution in the equilibrium U state, Y_N_ corresponds to the relative sum of amplitudes for the U-like distribution in the equilibrium N state, and Y_i_ denotes the relative sum of amplitudes for the U-like distribution at a given GdnHCl concentration. The relative sum of amplitudes was calculated as the sum of amplitudes for the U-like or N-like distribution divided by the total sum of amplitudes for both distributions. f_U_ derived from MEM-derived fluorescence lifetime distributions (Figures 4 and 5), had been shown previously to accurately estimate the relative fractions of N-like and U-like molecules present together.^13,92^

### Analysis of the fluorescence anisotropy decay traces

Time-resolved anisotropy decays were analyzed on the basis of the following equations.

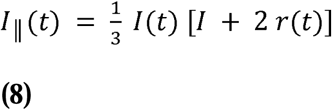

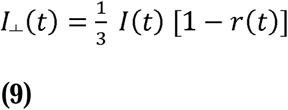

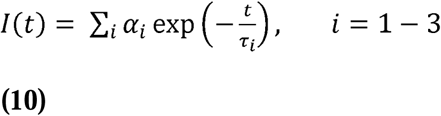

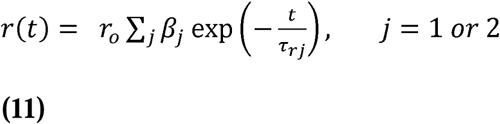

where I_‖_ and I_┴_ are the emission intensities collected at polarizations parallel or perpendicular to the polarization of the excitation beam. *I*(*t*) is the total fluorescence intensity at time *t*. r_o_ is the initial anisotropy, and α_i_and β_i_ are the amplitudes associated with the *i*th fluorescence lifetime and *j*th rotational correlation time such that [α_i_ = [β_j_= 1. In this model, each τ_i_is associated with both τ_r1_and τ_r2_. I_‖_ and I_┴_ were analyzed globally.^93^ The accuracy of the estimated anisotropy decay parameters depended significantly on the accuracy of the geometry (G) factor of the emission monochromator. Hence, extreme care was taken in the estimation of the G-factor as described previously.^21^ The steady-state fluorescence anisotropy (*r*_ss_) was calculated from the parameters obtained from the time domain data, using equation 12.

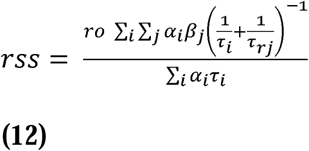

The measurement techniques and the data analysis procedures used were as described earlier.^21,77,94^

## ASSOCIATED CONTENT

### Supporting Information

SI contains supplementary figures.

## Acknowledgments

We thank Dr. Nilesh Aghera for generating the three Trp4 containing mutant variants of dcMN used in this study. We thank members of our laboratory and Dr MK Mathew for discussions. JBU is a National Science Chair of the Anusandhan National Research Foundation, Government of India. This work was funded by the Indian Institute of Science Education and Research Pune, and by the Anusandhan National Research Foundation.

## Accession codes

Double-chain monellin (dcMN), PDB entry 3MON; Single-chain monellin (MNEI), PDB entry 1IV7; UniProt entry P02881 and P02882 (MONA_DIOCU and MONB_DIOCU).

