## Supplementary Information for "Chain entropy modulates cooperativity selectively within intermediate sub-populations during protein unfolding"

### Supplementary Figures

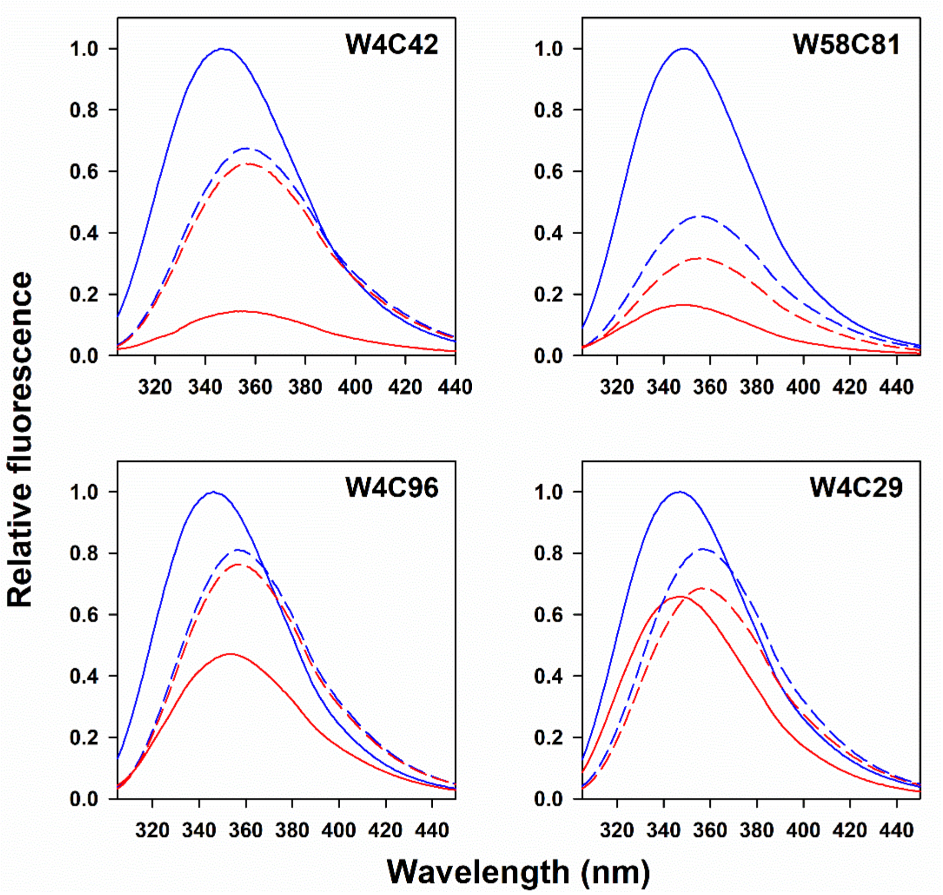


**Figure S1.** Fluorescence emission spectra of the different mutant variants of dcMN. The solid and dashed curves represent the spectra of the native and unfolded states (in 2 M GdnHCl), respectively, for the TNB-labeled (red) and unlabeled (blue) proteins. The spectra were normalized to the fluorescence value of the native unlabeled protein at 350 nm.

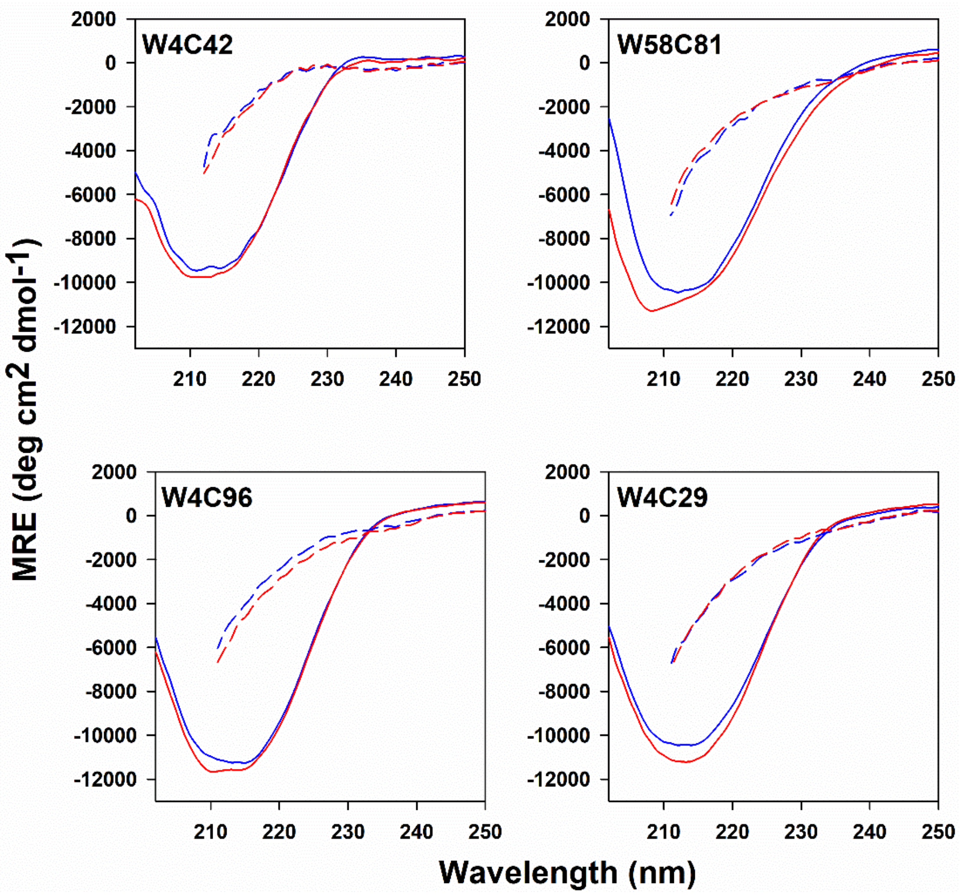


**Figure S2.** Far-UV CD spectra of the different mutant variants of dcMN. Spectra of the native (solid lines) and unfolded (dashed lines) states are shown for the TNB-labeled (red lines) and unlabeled (blue lines) proteins. Spectra of the unfolded proteins were measured in 2 M GdnHCl.

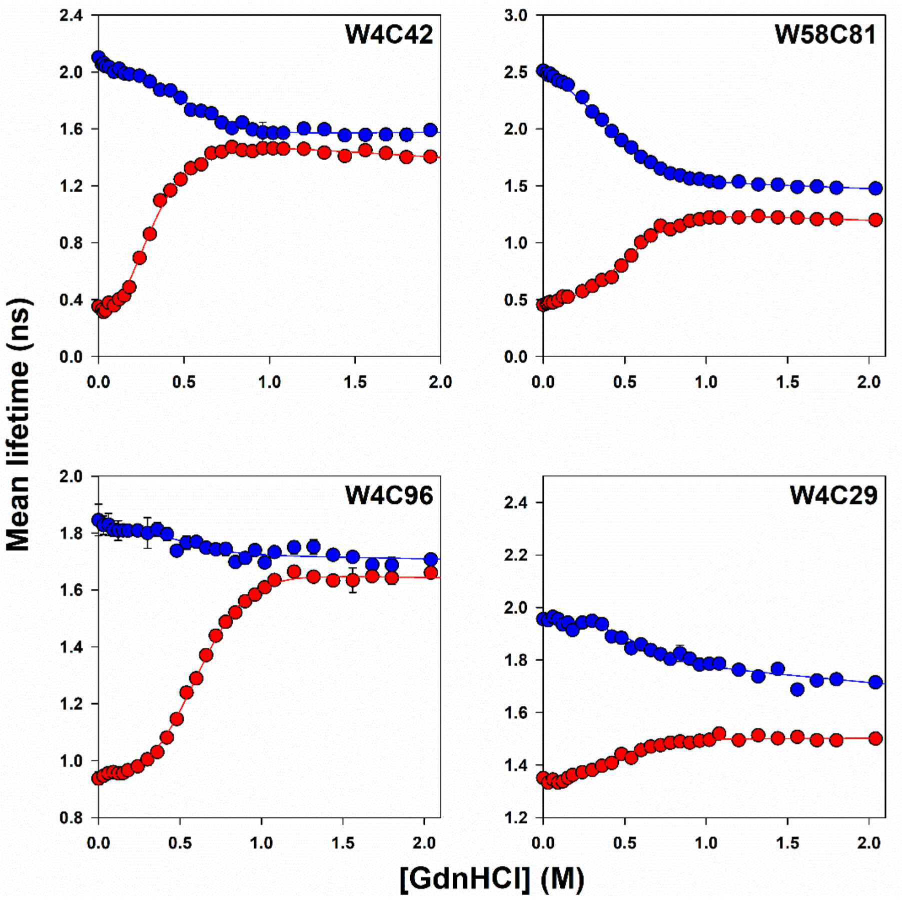


**Figure S3.** Equilibrium unfolding transitions of the different variants of dcMN monitored by time-resolved fluorescence measurements. Mean lifetimes were obtained from discrete analysis of the time-resolved fluorescence data. Blue and red circles correspond to the data obtained for the unlabeled and TNB-labeled proteins, respectively. The solid line passing through each dataset is a non-linear, least-squares fit to a two-state unfolding model for a heterodimeric protein.[1] The error bars represent the spread in the data obtained from two independent experiments.

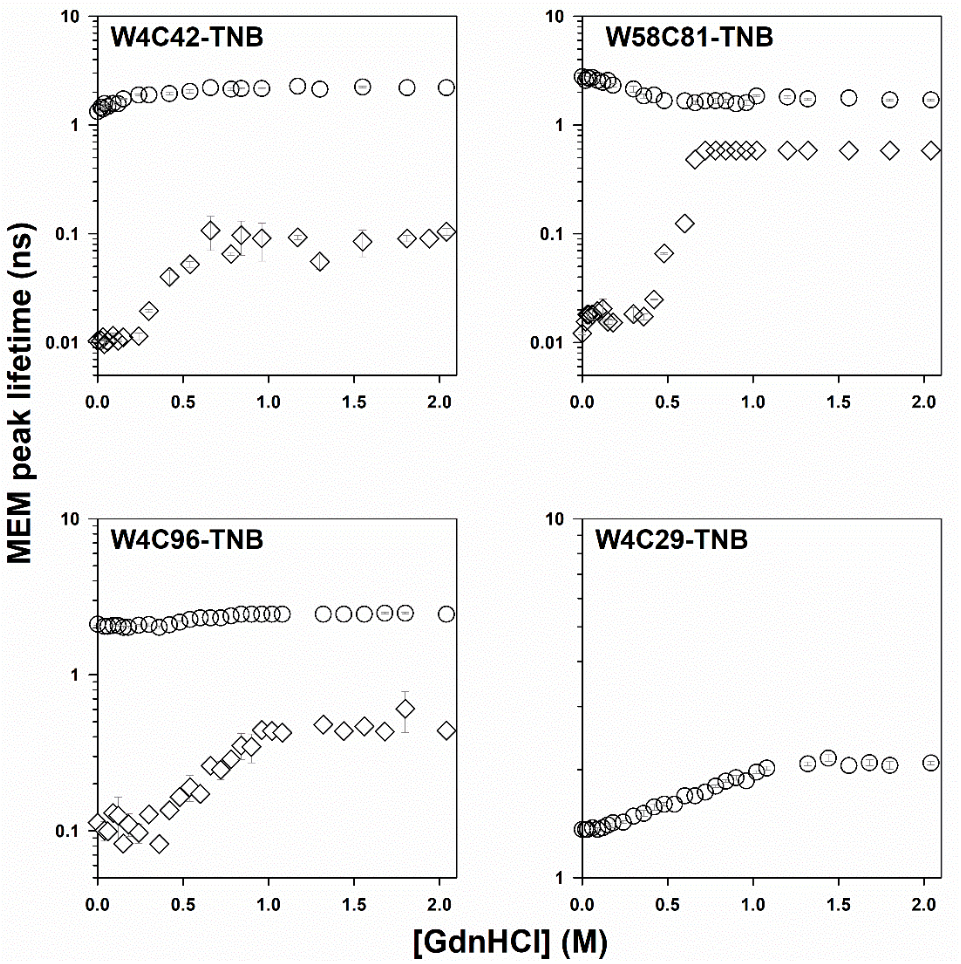


**Figure S4.** The shift in the MEM peak lifetime for the TNB-labeled variants of dcMN as a function of GdnHCl concentration. Diamonds and circles represent the lifetimes corresponding to the peaks in the MEM distributions arising from the N-like and U-like sub-populations, respectively. The circles in the panel for W4C29 correspond to the peak lifetimes of the observed unimodal fluorescence lifetime distributions. The error bars represent the spread in the data obtained from two independent experiments.

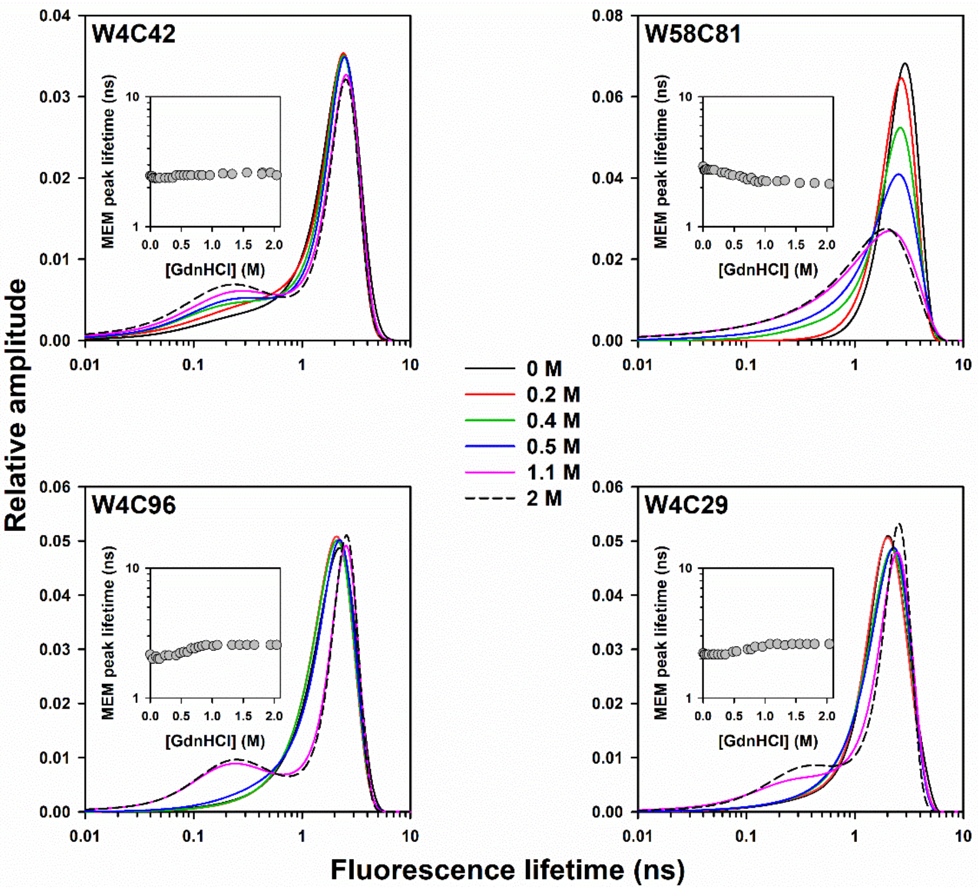


**Figure S5.** MEM-derived fluorescence lifetime distributions of the different mutant variants of dcMN at varying concentrations of GdnHCl. The colors of the curves represent the GdnHCl concentrations, as indicated. The inset in each panel shows the shift in the peak lifetime as a function of GdnHCl concentration. The error bars represent the spread in the data obtained from two independent experiments.

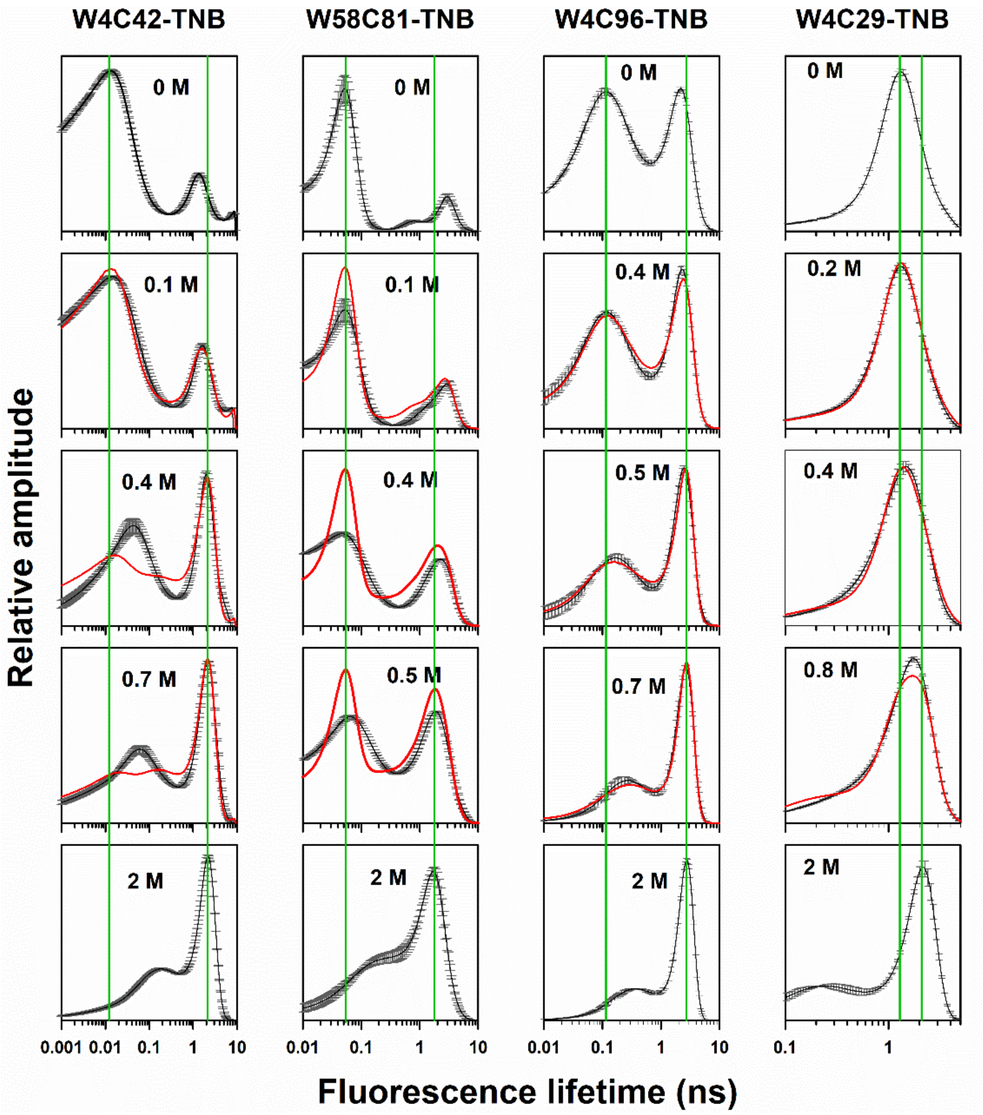


**Figure S6.** Fits of the fluorescence lifetime distributions obtained at different GdnHCl concentrations to a two-state N ↔ U model for equilibrium unfolding of the dcMN variants. The MEM-derived fluorescence lifetime distributions obtained at different GdnHCl concentrations (indicated in each panel) were fit to the weighted sum of the native state N(τ) and unfolded state U(τ) fluorescence lifetime distributions (equation 5, Materials and Methods). The black lines correspond to the experimentally determined distributions, and the red lines are fits to the two-state model. The error bars are the standard deviations of the MEM-derived fluorescence lifetimes obtained from multiple data acquisitions on the same sample. The top-most and bottom-most panels in each column show the fluorescence lifetime distributions used as the native N(τ) and unfolded U(τ) protein basis distributions, respectively. The vertical green lines indicate the peak lifetimes of the N-state and U-state fluorescence lifetime distributions. The x-axis has been plotted on a log scale, and the y-axis units are arbitrary.

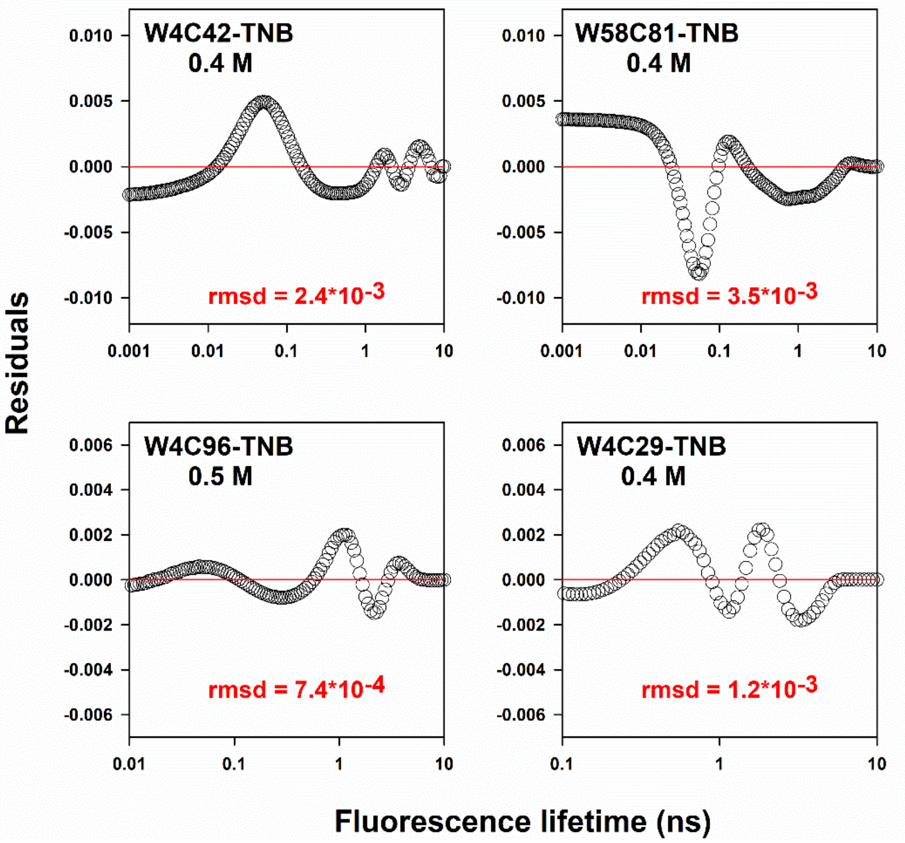


**Figure S7.** Residuals from the fits of the MEM-derived fluorescence lifetime distributions to the weighted sum of the mean of the MEM distributions determined for the N and U states (Figure S6). The residuals obtained for GdnHCl concentrations close to the mid-point of the unfolding transition are shown. Values in red correspond to the root mean square deviation (rmsd), which was obtained as the square root of the mean of the squares of residuals across all the lifetime values (x-axis).

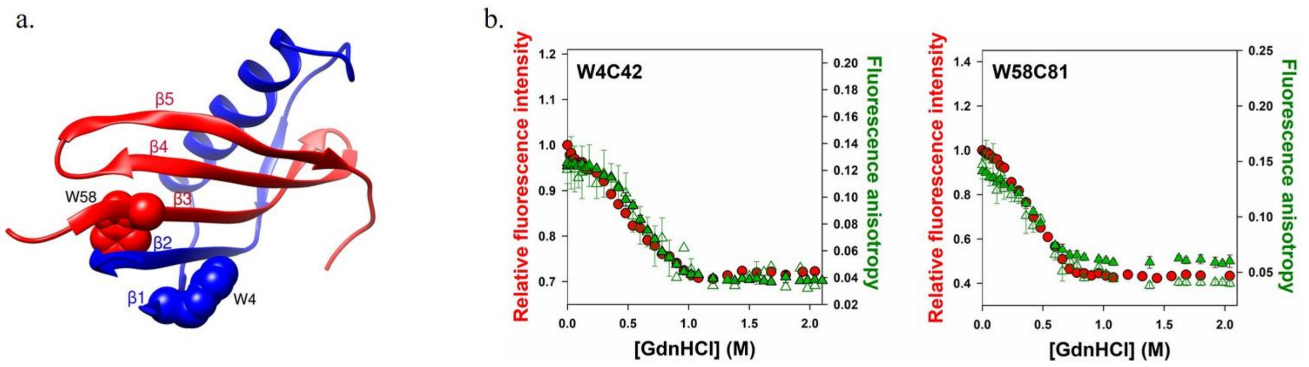


**Figure S8.** a) Structure of dcMN showing the locations of the two Trp residues. The side chains of Trp4 in chain B and Trp58 in chain A are shown in blue and red, respectively. b) Equilibrium unfolding transitions of the two variants were monitored by Trp fluorescence (red), steady-state anisotropy measured directly (filled green), and that derived from fits of time-resolved anisotropy decay traces (empty green). The fluorescence properties of Trp4 and Trp58 were monitored in W4C42 and W58C81, respectively.

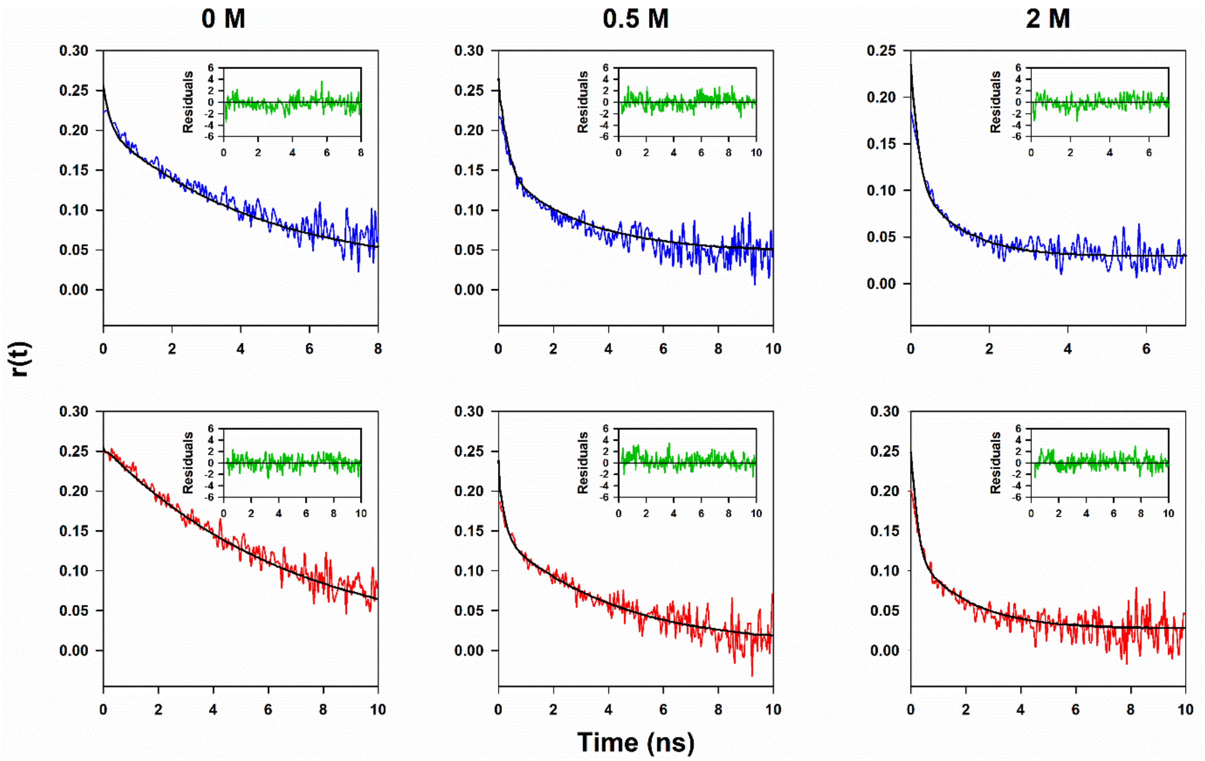


**Figure S9.** Time-resolved fluorescence anisotropy decay curves of Trp4 (blue) and Trp58 (red) monitored in W4C42 and W58C81, respectively, at the indicated concentrations of GdnHCl. The rotational correlation times and their corresponding relative amplitudes obtained from the fits (black solid lines) are shown in Figure 7. The decay curve of Trp58 in 0 M GdnHCl is the only one that fits well to a single-exponential equation. All other decay curves fit well to a two-exponential equation. Insets display the residuals of the fits.

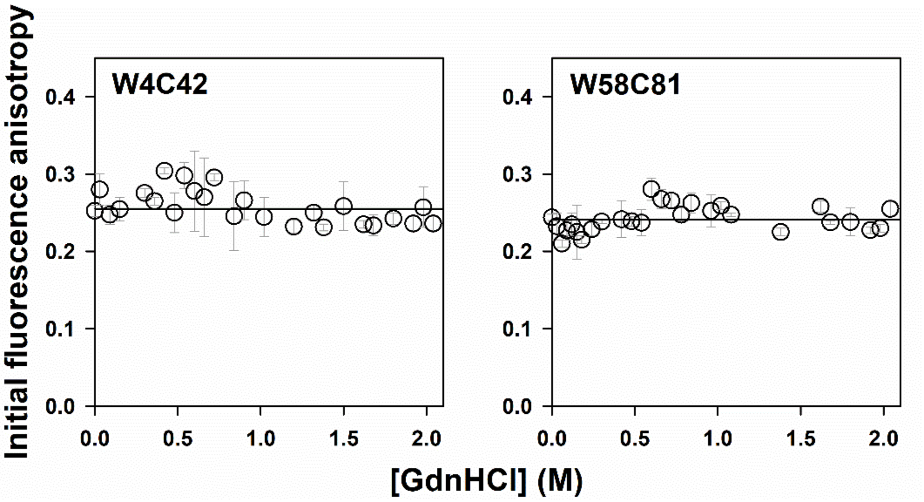


**Figure S10.** Equilibrium unfolding of W4C42 and W58C81 monitored by time-resolved fluorescence anisotropy. The dependences of the initial fluorescence anisotropy of the two variants on GdnHCl concentration are shown. The initial anisotropy of Trp4 was measured in W4C42 and that of Trp58 in W58C81. The error bars represent the spread in the data obtained from two independent experiments.

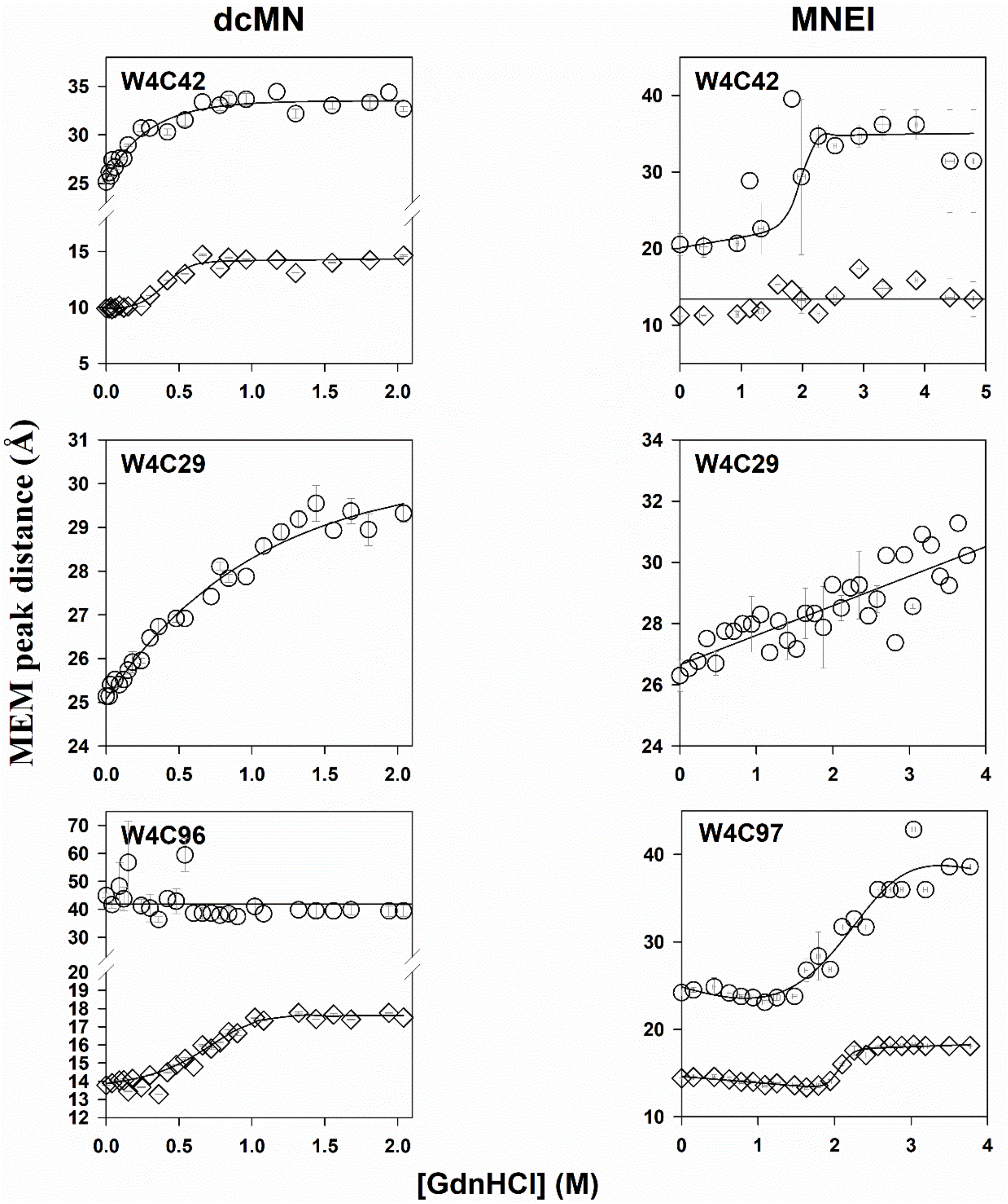


**Figure S11.** Comparison of the changes in the dimensions of the N-like and U-like sub-populations of dcMN and MNEI as a function of GdnHCl concentration. Symbol definitions are identical to those in Figure 6. The dcMN panels are reproduced from Figure 6, and the data for the corresponding segments of MNEI are reproduced from reference 2.

**Table S1.** Thermodynamic parameters obtained from fluorescence-monitored equilibrium unfolding measurements for the different mutant variants of dcMN at pH 8 and 25°C.

| Protein | Free energy of unfolding ($\Delta G_U$), kcal mol$^{-1}$ | Mid-point of unfolding ($C_m$), M |
|---|---|---|
| W4C42 | 11.0 ± 0.1 | 0.69 ± 0.04 |
| W4C42-TNB | 9.2 ± 0.4 | 0.27 ± 0.05 |
| W58C81 | 10.1 ± 0.1 | 0.46 ± 0.03 |
| W58C81-TNB | 9.6 ± 0.1 | 0.36 ± 0.04 |
| W4C96 | 10.6 ± 0.2 | 0.58 ± 0.05 |
| W4C96-TNB | 10.8 ± 0.1 | 0.64 ± 0.02 |
| W4C29 | 10.8 ± 0.4 | 0.63 ± 0.09 |
| W4C29-TNB | 10.5 ± 0.4 | 0.55 ± 0.08 |

**Table S2.** Energy transfer parameters of the native and unfolded states.

| Protein | FRET efficiency | Quantum yield | Overlap Integral, J*10^ (-13) | Förster distance, $R_o$ (Å) |
|---|---|---|---|---|
| W4C42_N | 0.84 | 0.1130 | 7.5 | 23.4 |
| W4C42_U | 0.10 | 0.0851 | 8.7 | 23.6 |
| W58C81_N | 0.82 | 0.1346 | 8.9 | 24.3 |
| W58C81_U | 0.18 | 0.0900 | 9.1 | 22.7 |
| W4C96_N | 0.50 | 0.0996 | 8.1 | 22.6 |
| W4C96_U | 0.02 | 0.0915 | 9.2 | 22.8 |
| W4C29_N | 0.31 | 0.1053 | 9.5 | 23.5 |
| W4C29_U | 0.12 | 0.0920 | 9.7 | 23.0 |
